# A role for brassinosteroid signaling in decision-making processes in the Arabidopsis seedling

**DOI:** 10.1101/2022.07.15.499689

**Authors:** Nils Kalbfuß, Alexander Strohmayr, Marcel Kegel, Lien Le, Friederike Grosseholz, Barbara Brunschweiger, Katharina Stöckl, Christian Wiese, Carina Franke, Caroline Schiestl, Sophia Prem, Shuyao Sha, Katrin Franz-Oberdorf, Juliane Hafermann, Marc Thiemé, Eva Facher, Wojciech Palubicki, Cordelia Bolle, Farhah F. Assaad

## Abstract

Plants often adapt to adverse conditions via differential growth, whereby limited resources are discriminately allocated to optimize the growth of one organ at the expense of another. Little is known about the decision-making processes that underly differential growth. In this study, we developed a screen to identify decision making mutants by deploying two tools that have been used in decision theory: a well-defined yet limited budget, as well as conflict-of-interest scenarios. A forward genetic screen that combined light and water withdrawal was carried out. This identified *BRASSINOSTEROID INSENSITIVE 2 (BIN2)* alleles as decision mutants with “confused” phenotypes. An assessment of organ and cell length suggested that hypocotyl elongation occurred predominantly via cellular elongation. In contrast, root growth appeared to be regulated by a combination of cell division and cell elongation or exit from the meristem. Brassinosteroid signalling mutants were most severely impaired in their ability to adjust cell geometry in the hypocotyl and cell elongation as a function of distance from the quiescent centre in the root tips. This study describes a novel paradigm for root growth under limiting conditions, which depends not only on hypocotyl-versus-root trade-offs in the allocation of limited resources, but also on an ability to deploy different strategies for root growth in response to multiple stress conditions.

## INTRODUCTION

Plants often adapt to adverse conditions via differential growth, whereby limited resources are differentially allocated to optimize the growth of one organ at the expense of another. A good example of this is the etiolation response (Arsovski et al., 2012), in which shoot growth is prioritized over root growth in the dark. Another example pertains to changes in root architecture in response to phosphate, nitrate or water deprivation (reviewed in Karban and Orrock 2018; Oyiga et al., 2020). Differential growth is also a key feature of plant responses to environmental stimulus when resources are sufficient, as seen in phototropism or gravitropism. In the case of phototropism, shoot curvature is achieved by differential growth within the stem, with one side growing faster than the other (Fankhauser und Christie 2015). In addition to differential growth decisions, plants need to assess trade-offs in the allocation of resources to defence versus growth (Leone et al., 2014; Lozano-Durán and Zipfel 2015; Campos et al., 2016; Fernández-Milmanda et al., 2020; Ortiz-Morea et al., 2020; van Butselaar and van den Ackerveken 2020). Plants also make choices as to when to initiate developmental processes or transitions such as germination, bud emergence, fruit set or leaf drop as well as the switch from vegetative to reproductive growth. The timing of floral transitions is impacted by environmental cues such as day length and temperature. Thus, milder winters are currently giving rise to earlier flowering in temperate climates. Furthermore, with changing climate the cues that guide decision making have become more erratic; in some cases, these cues even appear contradictory, as in the case of mild winters followed by late frosts or of drought followed by flooding. Understanding decision making in plants and how such processes respond to erratic or contradictory cues therefore becomes imperative to an understanding of the impact of changing climate.

In a judgement and decision-making model for plant behaviour, judgement is described as consisting of discrimination, assessment, recognition and categorization, whereas decision making involves an evaluation of the costs and benefits of alternative actions (Karban and Orrock 2018). Whether judgement and decision-making are empirically distinguishable remains to be determined. Nonetheless, it is interesting to reconsider the above-mentioned example of the floral transition within the framework of the judgement and decision-making model. When flowering occurs prematurely in a winter month, the questions that arise are what assumptions the plant can make as to how the spring will progress, as to when late frosts might set in, or as to the availability of water in the summer months to ensure a proper development of its fruit. It is not clear in this case what assumptions can be made and what degrees of uncertainty computed. Thus, in the formal language of decision theory (Bradley 2017), how a plant can assess the “state of the world” in the context of the floral transition appears unclear. The absence of such an assessment likely obscures the judgement required to inform decision-making. In brief, the floral transition might be classified as a choice made under considerable degrees of uncertainty. Simpler and clearer decision problems include the allocation of – or competition for – limiting resources (Shizgal 2012). Competition for light between neighbouring plants has, for example, been studied in *Potentilla reptans*, which was shown to adopt one of three strategies – vertical growth, shade tolerance or lateral avoidance – to optimize above ground responses to prevailing light-competition scenarios (Gruntman et al., 2017). Below ground, responses to variance in nutrient supply has been studied in split root pea exposed to variable and constant patches of soil under high or low mean nutrient concentrations; more roots developed in the variable patch when mean nutrients were low, whereas more roots developed in the constant patch when nutrients were high (Dener et al., 2016). This example depicts a clear assessment of risk and a consistently preferred outcome depending on mean nutrient levels: plants were risk adverse under high nutrient supply but risk prone when nutrients were low. While this example enables us to visualize decision-making in plants, the underlying regulatory networks remain poorly understood.

In this study, we explored a variety of screen conditions designed to best visualize decision-making in Arabidopsis. To identify major players, we performed a forward genetic screen. This calls for a simple decision problem. Therefore, we considered decisions reached under limiting conditions, as well as conflict of interest scenarios. We focused on the germinating seed, which has a limited energy budget clearly defined as the nutrient, oil and protein body reserves available in the Arabidopsis endosperm and embryo. Before the seed’s resources run out, the seedling must establish a root system capable of foraging for water and nutrients, and a photosynthetically active shoot system. Thus, the germinating seedling reaches binary shoot versus root growth decisions in terms of allocating the limited energy resources contained in the seed. We developed “conflict of interest” scenarios to monitor trade-offs between shoot versus root growth in the Arabidopsis seedling. These scenarios combine two abiotic stress factors that promote either hypocotyl or root growth. The ability to grow in response to limiting, adverse conditions is a survival strategy unique to plants; alternative responses adopted by yeast or animal cells include quiescence or the activation of apoptosis (Assaad, 2001). Our forward genetic screen identified BR signalling as playing a central role in decision-making in the seedling. We explore the strategies adopted to enable growth responses to abiotic stress cues within a limited budget, as well as the role of BR signaling in the deployment of such growth strategies.

## MATERIALS and METHODS

### Lines and growth conditions

*Arabidopsis thaliana* lines used in this study are listed in Table S1. Mutant lines were selected via the TAIR and NASC web sites (Swarbreck et al., 2008). EMS mutagenesis was carried out on Landsberg erecta (Ler) seed as described (Kim et al., 2006). Seed were surface sterilized, stratified at 4°C, and sown on Murashige and Skoog (MS) medium supplemented with B5 Vitamins (Merck group; https://www.sigmaaldrich.com). For nutrient stress conditions, NPK media was used. Plates were incubated under controlled growth chamber conditions (22 °C, 80 μmol m^-2^s^-1^). 10-day-old plate grown seedlings were used for organ length measurements and for scanning electron microscopy, and six or seven-day-old root tips were used for light microscopy. See supporting methods S1 and S2.

### Screen conditions

For all screen conditions, seed were germinated on nutrient or nutrient-deficient medium without a carbon source (see Fig. S1). Water deficit plates were prepared with PEG-6000 (Merck group; https://www.sigmaaldrich.com) as described in the supplemental methods. Plants were grown under optimal conditions (16: 8 hr light: dark photoperiod, 22°C, 180 μmol m^-2^s^-1^) at the TUMmesa ecotron (Roy et al., 2021) and the sterilized seed were imbibed at 4°C in the dark for 7 days (prior to plating) to break dormancy. Initial screen conditions compared growth on full-strength MS medium, with or without PEG-6000 for an additional −0.4 MPa. As the nutrients in MS medium generate a pressure of −0.2 MPa, initial screen conditions thus compared −0.2 MPa and −0.6 MPa. For optimized screen conditions, we later replaced full-strength MS by half-strength (½) MS medium, which corresponds to 0 MPa; thus, optimized screen conditions compare 0 MPa to −0.4 MPa. Under both initial and optimized screen conditions only the roots were exposed to the medium and, thus, to water stress; this was initially achieved via cutting a window in the agar and later via a plastic strip between the medium and the shoots. A final optimization was to incline the plates to promote root growth on the surface of the agar as opposed to in the agar. In the tables and graphs, we refer to initial versus optimized screen conditions to most accurately describe how the measurements were carried out. For growth under different light qualities blue, red or far-red LEDs (Quantum Devices) were employed. See supporting methods S3-S5.

### Molecular Methods

Mapping, positional cloning and allele sequencing were carried out as described in Jaber et al., 2010 using primers tabulated in the supporting method S6. qPCR was carried out as described in the supporting method S7.

### Light and electron microscopy

For scanning electron microscopy, a Zeiss (LEO) VP 438 microscope was operated at 15 kV. Fresh seedlings were placed onto stubs and examined immediately in low vacuum. Confocal microscopes used for imaging were an Olympus (www.olympus-ims.com) Fluoview 1000 confocal laser scanning microscope (CSLM) and a Leica (www.leica-microsystems.com) SP8 Hyvolution CSLM. FM4-64 staining was as described in Ravikumar et al., 2018. For GUS staining, micrographs were recorded with an Olympus BX61 microscope. See supporting methods S8 and S9.

### Image processing, data and statistical analysis

Shoot (hypocotyl) and root lengths were scored with the Image J free-hand tool (https://imagej.nih.gov). Images were processed with ImageJ, Adobe photoshop (www.adobe.com) and assembled with Adobe Illustrator. The hypocotyl volume was computed by assuming a cylindrical organ shape as follows: V = πWL, where W is the width and L the length of the hypocotyl.

Root and hypocotyl responses to water deficit in the dark (abbreviated as darkW) were computed as

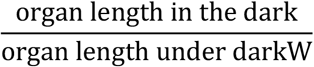

And for the ratio:

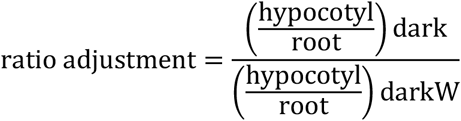

Due to variability between PEG lots and PEG plates (see text), we normalized each mutant to the corresponding wild-type ecotype (see Table S1) on the same plate. Thus, the normalized ratio adjustment to water stress in the dark (darkW) was computed as follows:

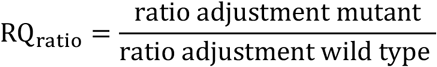

RQ_hypocotyl_ and RQ_root_ as well as light versus dark responses were computed in a similar fashion (see supplemental information). The mean RQ_ratio_ of at least three biological replicates (i.e. the seed stocks from different mother plants) is shown in Fig. 4 (see also supplemental information). Responses were considered to be attenuated for RQ_ratio_ < 0.8, normal in the 0.8-1.2 range, and exaggerated for RQ_ratio_ > 1.2. For volcano plots we plotted the mean RQ_ratio_ on the *X*-axis and the median *P_ratio_*-value on the *Y*-axis. *P*-values were computed with the Student’s *T*-test when the two populations had equal variances and with the Welch’s *T*-test when variances were unequal. That distributions were normal was verified with the Shapiro-Wilk test for selected samples, and in the rare cases where normality was not clear, we applied the non-parametric Mann-Whitney-U tests. The computations were carried out in excel for graphic rendition and verified in R. For multiple testing, as carried out when different wavelengths and light intensities were compared, we applied a Benjamini–Hochberg correction. This is specified in the legend where applicable. Responses were considered to be insignificant for *P*-values ≥ 0.05 and attenuated for *P*-values ≥ 0.00001. The median *P*-value for at least three replicates is shown in the volcano plots.

## RESULTS

### Hypocotyl growth in search of light is prioritized over root growth in search of nutrients in the young seedling

To understand decision making processes in the Arabidopsis seedling, we set up a “conflict of interest” scenario between shoot and root growth. To this end, seed were germinated under growth conditions designed to place contradictory demands on hypocotyl (growth at low light intensities or in the dark) or root growth (nutrient deficiency or water stress). We first tested several nutrient media for their ability to promote trade-offs between hypocotyl versus primary root growth in dark-grown seedlings; these include -P, -N, and -K media lacking phosphate, nitrate, or potassium (Fig. S1). We also tested low levels of osmotic stress (100 mM mannitol; Fig. S2) as well as salt stress (50-100 mM NaCl; Fig. S2), which has been shown to impair hypocotyl elongation in response to far-red light (Hayes et al., 2019). None of these media, however, gave rise to considerable or reproducible trade-offs between root and hypocotyl growth, by which we refer to the growth of one organ at the expense of another (Fig. S1; Fig. S2). There was a clear priority for hypocotyl growth in search of light over primary root growth in search of nutrients (Fig. S1d). It is to be noted in this context that we are looking not at root architecture but exclusively at primary root growth and that the seed and embryo contain sufficient nutrients to support the initial phases of seedling growth (Lott and West 2001).

### Opposing gradients of water stress and light intensity enable us to visualize shoot versus root growth trade-offs in the seedling

We next attempted to identify what resources might be as important to a germinating seedling as the light. Here, we explored the relative importance of water. To this end, we germinated seedlings in the presence of water stress, using polyethylene glycol (PEG) to withdraw available water from the medium in a standardized tissue culture setting (Michel and Kaufmann 1973; van der Weele et al., 2000). In the light, PEG significantly decreased hypocotyl growth (*P_hypocotyl_* = 3E^-48^), but this was not mirrored by an increase in root growth; root growth was, in fact, slightly reduced (*P_root_* = 1.3E^-02^; Fig. S3a). In the dark, increasing degrees of water stress considerably and reproducibly increased root length and decreased hypocotyl length (three-fold change for the hypocotyl and almost four-fold change for the root for −0.7 MPa compared to −0.2 MPa; Fig. 1 a-e; Fig. S3b-d). In the light, water stress decreased the total length of the seedlings (*P_total_* = 8.2E^-4^; Fig. S3a). In the dark, however, water stress did not impact the total length of the seedling (for ≤ −0.6 MPa compared to −0.2; Fig. 1e cf Fig. S3a). In comparison to water-stress in the light, we were clearly observing tradeoffs between hypocotyl and root growth in response to a gradient of water stress in dark-grown seedlings. We then tested a gradient of decreasing light intensity. Under low light, the tradeoff was more pronounced, with longer roots and shorter hypocotyls than in the dark, even when the light intensity was as low as 2 μmol m^-2^ s^-1^ (Fig. 1f-i). Similarly, for both blue and red light, hypocotyl length increased whereas root length decreased with decreasing light intensity and this trend was enhanced by water stress (−0.6 MPa; Fig. S4a, S4b). Far-red light showed a comparable trend (Fig. S4c). In conclusion, using opposite gradients of decreasing light intensity versus increasing water stress, we were able to fine tune root growth at the expense of hypocotyl growth.

**Figure 1.**
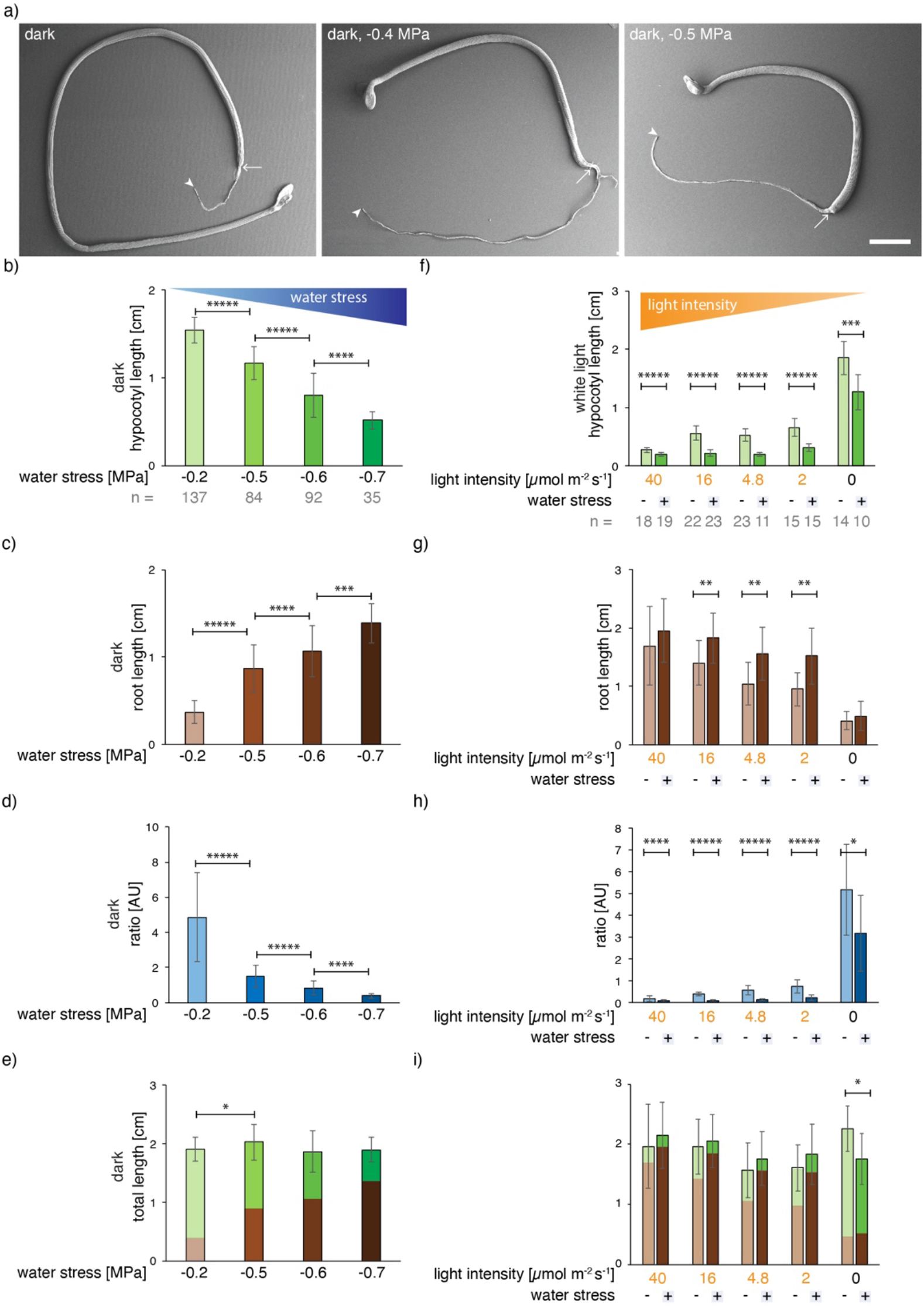
Hypocotyl versus root growth in response to light and water availability. Wild type (Col-0). (a-e) germination in the dark on a gradient of water stress ranging from 0 to −0.7 MPa. Recordings were taken 10 days after germination. (a) scanning electron micrographs; arrows point to the hypocotyl/root junction and arrowheads to the end of the root. (b-e) panels are from the same experiment. Water stress applied in the dark increases root length (c) at the expense of hypocotyl length (b), giving rise to a decrease in the hypocotyl/root ratio (d). Note that the total seedling length does not vary at −0.2 MPa versus −0.6 or – 0.7 MPa (e). Thus, there was a clear tradeoff, by which we refer to the growth of one organ at the expense of another. (f-i) germination on MS medium under white light at varying intensities ranging from 40 to 0 μmol m^-2^ s^-1^, with or without −0.6 MPa water stress. A decreasing light intensity gradient increases hypocotyl length (f) at the expense of root length (g). The hypocotyl/root ratio was calculated (h). The number (n) of seedlings measured per condition is in grey below the mean ±StDev bar graphs. *P*-values were computed with a two-tailed student’s *T*-test and are represented as follows: *: 0.05 – 0.01; **: 0.01 – 0.001; ***: 0.001 – 0.0001; ****: 0.0001 – 0.00001, *****: < 0.00001. See related Fig. S1, S2, S3, S4 and S5.

We next asked whether the reduction in hypocotyl length observed in response to water stress applied in the dark was accompanied by a change in organ width and, ultimately, volume. To this end, seedlings were imaged by scanning electron microscopy (Fig. 1a) and the lengths and widths of the hypocotyls measured (Fig. S5a, S5b). In a comparison between light to dark, the hypocotyl length and volume increased despite a decrease in hypocotyl width (Fig. S5a-c). When water stress was applied to dark-germinated seedlings, the reduced length of the hypocotyl was not accompanied by increase in width; in fact, a slight reduction in width was observed and overall the volume of the hypocotyl was two-fold reduced (Fig. S5a-c). The change in organ volume suggests that resources such as water, which accounts for organ volume to a large extent, are being differentially allocated in response to light and water stress. In summary, hypocotyl and root lengths and the ratio thereof are exquisitely fine-tuned to the wavelength and intensity of the light source, and to the severity of water stress. In the case of the seedling, what we observe is a clear consistency in a continuum of preferred outcomes along a gradient in response to opposing gradients of light intensity and water stress.

### A forward genetic screen for “decision” mutants identifies BRASSINOSTEROID INSENSITIVE 2 (BIN2)

The fine-tuning of hypocotyl/root ratios can conceptually be broken down into four steps: (i) sensing, (ii) downstream signaling, (iii) decision making processes, and (iv) the execution of these decisions (the action). The gradients of hypocotyl and root growth we describe in response to our screen conditions can help distinguish between mutants impaired in the process of decision making *per se,* versus mutants with primary defects in one of the other steps (sensing, signaling or execution). Perception mutants would fail to perceive light or water stress; a good example of this is the *cry1 cry2 phyA phyB* quadruple photoreceptor mutant (Mazzella and Casal 2001), which had a severely impaired light response (Fig. S4f), but a “normal” response to water stress in the dark (Fig. S4g). In contrast, execution mutants may have aberrantly short hypocotyls or roots that are nonetheless capable of differentially increasing in length depending on the stress conditions. Decision mutants would differ from perception or execution mutants as they would clearly perceive the stress factors yet fail to adequately adjust their hypocotyl/root ratios in response to a gradient of multiple stress conditions. Failure to adjust organ lengths would be seen as a non-significant response, or as a significant response but in the wrong direction as compared to the wild type. We thus used organ lengths, the hypocotyl/root ratio and the significance of the responses as decision read outs. We specifically looked for mutants in which at least one organ exceeded wild-type length under darkW. To operate under limiting conditions, we germinated seedlings in the dark on media lacking a carbon source (see Fig. S1), avoiding even low levels of light.

To identify genes implicated in decision making, we performed a genetic screen in two consecutive steps (Fig. 2a). In the first step, we germinated seed in the dark. In the second round, viable mutants with aberrant hypocotyl to root ratios in round 1 were rescreened for their ability to adjust their hypocotyl to root ratios in response to water stress in the dark. We initially screened in the dark because the high variance in root growth under water deficit in the dark in the wild-type (see below) would obscure the distinction between putative mutants versus stochastically occurring wild-type seedlings with short roots under darkW. 83000 EMS-mutagenized M2 seed were subjected to the first screen and over 100 viable mutants rescreened in the dark in the presence or absence of water stress. 19 viable seedlings with aberrant hypocotyl to root ratios in our forward genetic screen had brassinosteroid-related dwarf phenotypes (Fig. 2b; Fig. S6a, S6c). Of these, one – named B1 – had a very pronounced inability to adjust its hypocotyl/root ratio in response to multiple stress conditions: while the root response to water stress in the dark was only slightly attenuated (*P_root_* = 6E^-05^; Fig. 2c), the hypocotyl response was aberrant (*P_hypocotyl_* = 0.05; Fig. 2c) and the ratio adjustment insignificant (*P_ratio_* = 0.83; Fig. 2c). Positional cloning and allele sequencing suggested that the B1 locus encoded BRASSINOSTEROID-INSENSTIVE 2 (BIN2; Fig. 2d, 2e). Our B1 allele carries a TREE domain E263K mutation, identical to *bin2-1* and shown to block the BR-induced ubiquitination and degradation of BIN2 (Peng et al., 2008). Segregation analysis showed that our B1 *bin2* allele was semi dominant, like all TREE domain mutations of *bin2-1.* We sequenced 19 F2 individuals of the segregating mapping population and found that the E263K point mutation absolutely segregated with the phenotype (Table S2). For complementation analysis, we crossed our B1 mutant with a known *bin2-1* allele (Li et al., 2001); F1 individuals had phenotypes on soil that were characteristic of homozygous *bin2* plants and, upon sequencing, exhibited EMS-induced G to A transitions at position 989 for both the B1 mutation and the *bin2-1* allele (Fig. 2e). We complemented B1 with *bin2-1* and *ucu1* alleles and compared it to *bin2-1, ucu1* and *dwarf12* (Pérez-Pérez et al., 2002; Choe et al., 2002) alleles at the BIN2 locus; these three published mutant lines exhibited the same behavior as B1, including semi-dominance and partial etiolation. Indeed, the B1 phenotype was similar to *bin2-1* under our multiple stress conditions (Fig. S6b; compare B1 in Fig. 2c to *bin2-1* in Fig. 3d). In conclusion, positional cloning, allele sequencing, segregation analysis, complementation analysis and phenotypic analyses show that the B1 locus encodes BIN2.

**Figure 2.**
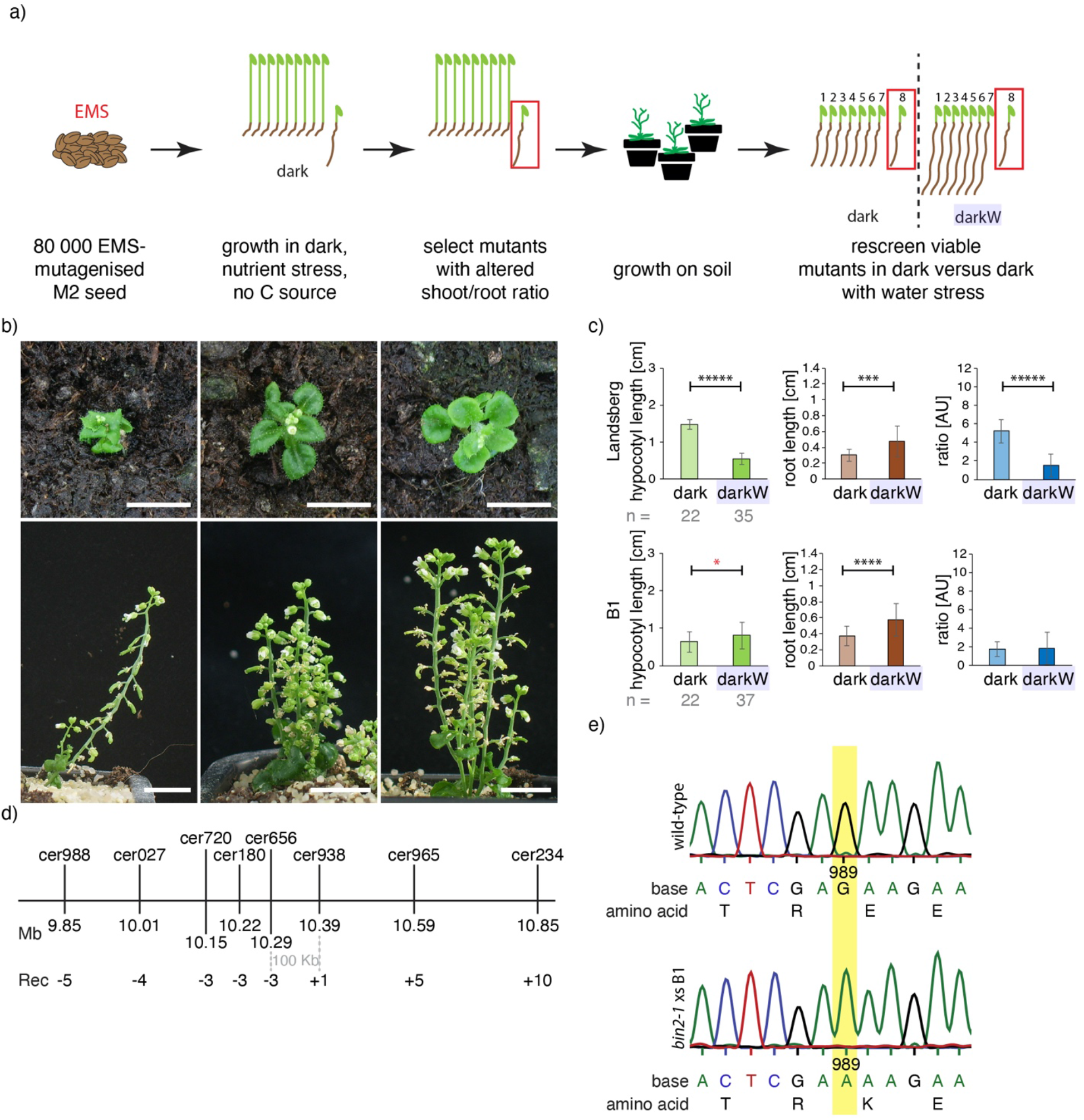
Identification of BIN2 and its role in hypocotyl/root tradeoffs. (a) A screen for decision mutants; two consecutive steps are depicted (see text). (b) Dwarf mutants with curled-in leaves, a BR-related phenotype, identified in our forward genetic screen on the basis of altered hypocotyl/root ratios in the dark. Scale bars = 1cm. (c) B1 mutants fail to adjust their hypocotyl/root ratios in response to water stress in the dark (darkW). Note that hypocotyls had an inverse response (significance marked with red asterisk). The number (n) of seedlings measured per condition is in grey below mean ±StDev bar graphs. *P*-values were computed with a two-tailed student’s *T*-test and are represented as follows: *: 0.05 – 0.01; ***: 0.001 – 0.0001; ****: 0.0001 – 0.00001, *****: < 0.00001. (d) Fine mapping of mutant B1 identifies a 100 kb interval on chromosome 4, which spans the *BIN2* locus. Markers used for mapping are depicted in abbreviated form above the line and in full detail in the supplemental methods. Rec: recombinants. See supporting methods M6 for mapping and Table S2 for segregation analyses. (e) Sequencing of the BIN2 TREE domain in F1 segregants of a complementation cross between *bin2-1* and B1 shows that B1 harbors a G to A transition at position 989 (yellow highlight), giving rise to an E263K mutation identical to that of *bin2-1.* n = 10 F1 plants were sequenced. See related Fig. S6.

### The BR pathway is implicated in hypocotyl versus root tradeoffs in the Arabidopsis seedling

Brassinosteroids are known to be involved in the response to abiotic stress cues such as drought and salinity (Fàbregas et al., 2018; Cui et al., 2019; Hayes et al., 2019). To analyse the role of BR signalling in decision making processes, we studied a set of known mutants impaired in BR biosynthesis, perception, signalling or in BR-responsive gene expression (Table S1). In addition to bar graphs representing hypocotyl and root lengths (Fig. 3), the distribution of datapoints was represented by violin plots (Fig. 4a-c, Fig. S7). The violin plots compare organ length distributions in mutants versus the corresponding wild-type ecotype, which depicts dwarfism in some brassinosteroid mutants. It is also apparent that wild-type (Col-0) root length varies under water-deficit in the dark (Fig. S7). Although we have optimized protocols for PEG plates to the best of our ability, there is still a lot-to-lot and plate-to-plate variation. This emphasizes the need for normalizing each mutant line to its corresponding wild-type ecotype on the same (PEG) plate in the same experiment. To this end, the response to water stress in the dark was represented as a normalized response quotient (RQ), which is an indication of how much the mutant deviates from the corresponding wild type (Fig. 4e; see methods). We used the normalized ratio response RQ_ratio_ as our main “decision” readout (Fig. 4e), and show RQ_hypocotyl_ and RQ_root_ in the supplement (Fig. S8). This is because water stress in the dark is a “conflict of interest” scenario in which hypocotyl and root growth have competing interests (see Fig. 1). We reason that mutants unable to integrate environmental cues might have a “confused phenotype” under our multiple stress conditions. The read out for a “confused phenotype” would translate into erratic (i.e. highly variable) hypocotyl versus root lengths. This high variance would, in turn, translate into a low signal to noise ratio, and this can be seen as a high *P*-value. We, therefore, plotted the median *P*-values against the normalized response quotients (referred to as volcano plots; mean RQ_ratio_ in Fig. 4f). Mutants with high *P*-values and low response quotients would be considered “confused” and these would map in the lower left quadrant of the RQ_ratio_ volcano plot (response to water stress in the dark, Fig. 4f).

**Figure 3.**
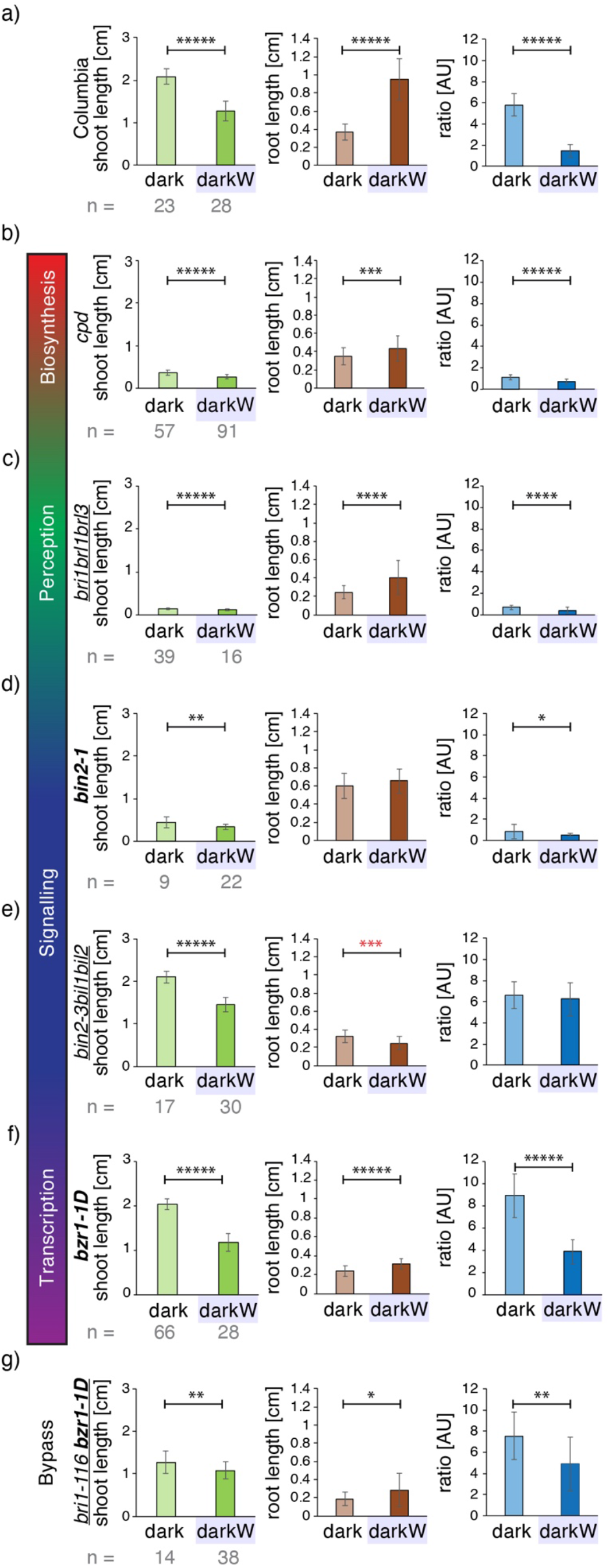
Role of BR signaling in hypocotyl/root tradeoffs. Seedlings were germinated on ½ MS in the dark (dark) or in dark with −0.4MPa water stress (darkW). (a) Col-0 (wild type). (b) BR biosynthesis mutant *cpd.* (c) BR perception mutant *bri1brl1brl3* (a segregating triple null). (d) BR signaling mutant *bin2-1* (a semidominant gain of function allele). (e) *bin2-3bil1bil2* tripple knockout; note that, in stark contrast to the wild type, the *bin2-3bil1bil2* tripple knock-out has shorter roots under darkW than in the dark (red asterisks). (f) Transcription factor mutant *bzr1-1D,* a dominant allele. (g) BIN2 bypass mutant *bri-116 bzr1-1D.* Note that *bin2-1* mutants have a severely attenuated hypocotyl response and no root response (d), *bin2-3bil1bil2* mutants have an inverse root response (red asterisks denote a significant response in the opposite direction to the wild type) and no ratio response (e), and *bri-116 bzr1-1D* severely attenuated hypocotyl, root and ratio responses (g). Null alleles are depicted in regular font, semi-dominant or dominant in bold and higher order mutants are underlined. At least 3 experiments were performed for each line, and a representative one is shown here on the basis of RQ and P values (see Fig. 4e,f). The number (n) of seedlings measured per condition is in grey below the mean ±StDev bar graphs. *P*-values were computed with a two-tailed student’s *T*-test and are represented as follows: *: 0.05 – 0.01; **: 0.01 – 0.001; ***: 0.001 – 0.0001; ****: 0.0001 – 0.00001, *****: < 0.00001. For mean RQ values and median *P*-values see Fig. 4e, f, and related Fig. S8. Mutant alleles and the corresponding ecotypes are described in Table S1. See related Fig. S6, S7, S8, S9, S10, S11.

**Figure 4.**
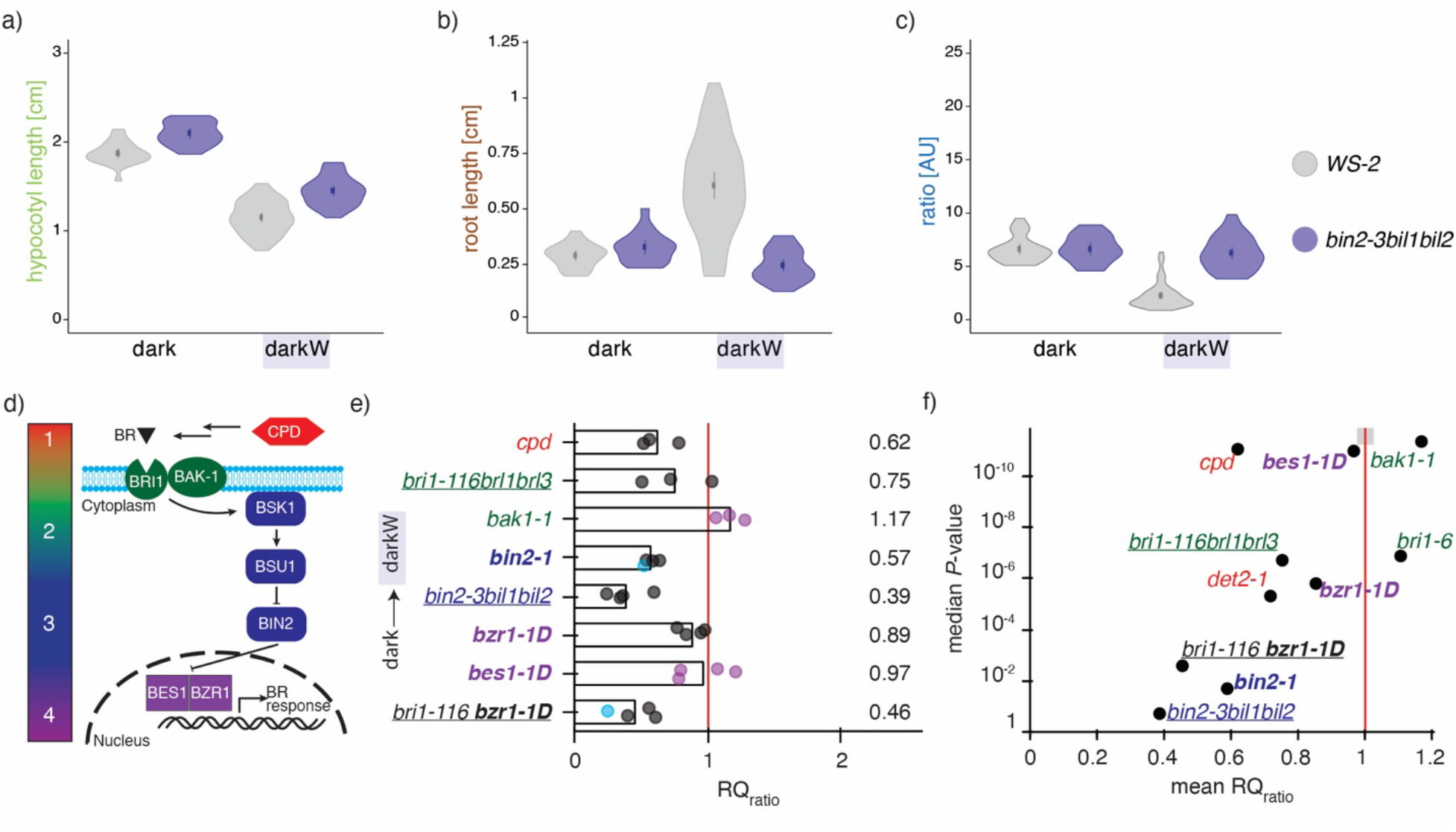
Responses of BR pathway mutants to water stress in the dark: violin plots, response quotients and volcano plots. (a-c) violin plots of the hypocotyl (a), root (b) and ratio responses (c) of the triple *bin2-3bil1bil2* knock out line shown in Fig. 3e, with the wild-type ecotype (Ws-2) as reference. The dot represents the mean and the line the 95% confidence interval. Note the high variance of the wild-type root response under darkW. *bin2-3bil1bil2* mutants qualified as decision mutants on 3 counts: (i) failure to adjust the hypocotyl/root ratio to darkW (the ratio for darkW is the same as for dark in panel c), (ii) low or non-significant P-value (see panel f below) and (iii) one organ (here the hypocotyl in panel a) exceeded wild-type length under darkW. (d) Color coding of steps in the BR signalling pathway; 1: biosynthesis, 2: perception, 3: signalling, 4: transcription. See text for further detail. (e) RQ_ratio_ response quotient of the hypocotyl/root ratios under dark/darkW conditions, normalized to the wild-type ratio quotient; a value of 1 (vertical red line) indicates that the response to a shift from dark to darkW is similar to that of the respective wild-type ecotype. Each replicate is represented by a dot; purple dots are for initial and grey dots for optimized screen conditions; blue dots are for data from SEM measurements. Note that the triple *bin2bil1bil2* knock out has the strongest phenotype, followed by *bri1-116 bzr1-1D* and *bin2-1.* (f) Volcano plot with the mean RQ_ratio_ depicted in (e) on the *X*-axis and the median *P*-Value of the response on the *Y*-axis (negative log scale; a median of all replicates was used). Mutants in the lower left quadrant are considered to have a “confused decision phenotype” (see text). Only *bin2* alleles or bypass mutants map to this quadrant. The area shaded in grey on the red line is where Col-0 would theoretically map onto the plot. Mutant alleles and their ecotypes are described in Table S1. Null alleles are depicted in regular font, semi-dominant or dominant in bold and higher order mutants are underlined. See related Fig. S6, S7, S8, S9, S10 and S11.

In the BR pathway (Fig. 4d), we first looked at *cpd*, a BR biosynthesis mutant. *CPD (CONSTITUTIVE PHOTOMORPHOGENIC DWARF)* encodes a cytochrome P450 monooxygenase involved in the C6 oxidation pathway of brassinolide biosynthesis (Szekeres et al., 1996). *cpd* mutants are dwarfs with attenuated but significant responses to water stress in the dark (Fig. 3b; Fig. S7; Fig. 4e, 4f; Fig. S8). We then turned to higher order mutants of the BR receptor, BRI1, and its homologues, *bri1-116 brl1 brl3* (Kang et al., 2017). *bri1-116 brl1 brl3* mutants had an attenuated response to water stress in the dark; however, as for *cpd*, all trends were similar to the wild type and all responses significant (Fig. 3c; Fig. S7; Fig. 4e, f; Fig. S8). An allele of the BAK1 coreceptor, *bak1-1* (Li et al., 2002), showed fairly similar responses to a hypomorphic *BRI1* allele, *bri1-6* (Fig. 4f).

When BR binds to the extracellular domain of the BRI1 receptor, the BIN2 BRASSIONSTEROID INSENSISTIVE 2 shaggy-like kinase is inactivated (Fig. 4d). We assessed the phenotype of the semi-dominant *bin2-1* allele, which has an identical amino acid change (TREE to TRKE) in the tree domain as our *bin2* B1 allele. For *bin2-1,* the response to water stress in the dark was severely impaired: the hypocotyl and root responses were non-significant (median *P*_hypocotyl_ = 0.11; median *P*_root_ = 0.07; Fig. 3d; Fig. S7; Fig. S8), and the hypocotyl/root ratio response considerably attenuated (mean RQ_ratio_ = 0.57; median P_ratio_ = 0.02; Fig. 4e, 4f). *bin2-1* seedlings are dwarfs with severe and pleiotropic phenotypes (Li et al., 2001), including a reduction in seed size. We, therefore, turned to higher order null alleles. The Arabidopsis genome encodes ten shaggy-like kinases, of which three in the BIN2 clade have been shown to function redundantly (Yan et al., 2009). A triple knock-out of all three clade 2 shaggy like kinases, *bin2-3bil1bil2*, has a large stature, short roots and an overall phenotype characteristic of plants with enhanced BR signaling outputs (Yan et al., 2009). Seed weight in *bin2-3bil1bil2* did not differ from the wild type (Fig. S9a). This mutant was severely impacted, with a non-significant (median *P_root_* = 0.44; mean RQ_root_ = 2.42; Fig. S8b) or aberrant root response to water stress applied in the dark (Fig. 3e; Fig. 4e, f). *bin2-3bil1bil2* mutants fit the above definition of decision mutants as they have a significant root response but in the wrong direction as compared to the wild type, as denoted by red asterisks (Fig. 3e). The hypocotyl/root ratio response was also non-significant (median *P_ratio_* = 0.19) and severely impaired (mean RQ_ratio_ = 0.39; Fig. 3e; Fig. 4e, f). Thus, both semi-dominant and recessive mutations in clade 2 shaggy-like kinases impair differential growth responses to light and water stress in the seedling.

The active BIN2 kinase phosphorylates BZR and BES transcription factors, which reduces their DNA-binding activity, excludes them from the nucleus and targets them for degradation (Nolan et al., 2020). By inactivating BIN2, BR signaling results in the accumulation of unphosphorylated BZR1/2 in the nucleus and a concomitant expression of BR-target genes. We assessed the behavior of a dominant BZR1 allele, *bzr1-1D* (Wang et al., 2002), which is, like *bin2-3bil1bil2,* characterized by enhanced BR signaling outputs (Yan et al., 2009). Seed weight was not impacted in *bzr1-1D* (Fig. S9b). With respect to water stress in the dark, the root and hypocotyl/root ratio responses were significant and fairly similar to the wild type (mean RQ_root_ = 1.17; mean RQ_ratio_ = 0.89: Fig. 3f; Fig. 4e, 4f; Fig. S8b). A BES1 dominant allele, *bes1-1D* (Yin et al., 2002), had a weaker phenotype than *bzr1-1D* in that it did not exhibit a considerable difference from the wild type in terms of its response to water stress in the dark (mean RQ_ratio_ = 0.97; median *P_ratio_* = 2E^-11^; Fig. 4e, f).

Looking at BR pathway mutants suggests that BR signaling is implicated in hypocotyl versus root growth tradeoffs in the Arabidopsis seedling. To address this hypothesis, we looked at the *bri1-116 bzr1-1D* double mutant (Wang et al., 2002), in which a null non-viable BRI1 receptor mutant is partially rescued by a dominant BZR1 transcription factor allele. In this line, the BR pathway, including BIN2 function within this pathway, is effectively bypassed. *bri1-116 bzr1-1D* double mutants had a severely attenuated hypocotyl response to water stress in the dark (mean RQ_hypocotyl_ = 0.58; median *P_hypocotyl_* = 0.01; Fig. 3g; Fig. S8). Similarly, the double mutants had very short roots in the dark that elongated erratically (RQ_root_ = 1.43; non-significant median *P_root_* = 0.07; note the large variance in Fig. 3g; Fig. S8b) when water stress was applied in the dark. The hypocotyl to root ratio adjustment was severely impaired (mean RQ_ratio_ = 0.46; median *P_ratio_* = 0.003; Fig. 3g; Fig. 4e, f). In summary, BR bypass mutants mapped together with *bin2* gain of function and loss of function mutants in the same quadrant of the volcano plots, showing a “confused” decision phenotype (high *P*-value, low ratio quotient; Fig. 4f). We conclude that BR signaling is required for differential growth decisions in the Arabidopsis seedling.

### BR pathway mutants perceive light and water withdrawal

We next asked whether BR mutants are capable of perceiving and responding to light or waterstress. We first compared light versus dark conditions. Some BR pathway mutants exhibited etiolated phenotypes (short hypocotyl, long root, at least partially open cotyledons in the dark), as described in the literature (Szekeres et al., 1996). Nonetheless, and even though responses were attenuated in some mutants, all BR pathway mutants had significant responses to light (shorter hypocotyls, longer roots as compared to dark-grown seedlings; Fig. S10; Fig. S11). *bin2-1* had a dwarf phenotype and an aberrant hypocotyl/root ratio, with a short hypocotyl and a relatively long root even in the dark (Fig. S10d; Fig. S11). Interestingly, the BR mutant lines with the strongest etiolation phenotypes *(cpd* and *bri1-116brl1brl3*, Fig. S11a,b) in the dark were not the ones with the strongest deviation from the wild type under water deficit in the dark (Fig. S8). We also looked at the expression levels of the light responsive gene *LHCB1.2* via qPCR in wild-type Ws-2 versus *bin2-3bil1bil2.* The data show that *LHCB1.2* gene expression is light-regulated in *bin2-3bil1bil2* seedlings (Fig. S12). We further looked at the response of selected BR mutants to water stress (−0.4MPa) in the light. Here, we focused on the hypocotyl response as this was clear and consistent in the wild type (Fig. S3a). We found that both *bin2-3bil1bil2* and *bzr1-1D* mutants were unimpaired in their hypocotyl responses to water stress in the light (Fig. S13). We conclude that the investigated BR mutants are not primarily impaired in their ability to perceive light or water stress.

### BR signalling is required for a differential regulation of cell anisotropy in the hypocotyl

Hypocotyl elongation in the dark is known to occur via cellular elongation, with no significant contribution of cell division in the epidermis or cortex (Gendreau et al., 1997). An assessment of organ and cell length in this study corroborated these findings, also for water stress in the dark. Indeed, we show that cellular parameters (cell length and width, assessed in scanning electron micrographs of hypocotyl cells) accounted for more than 73% of the observed differences in organ length and >89% of the differences in organ width (see Fig. 5a, b for cellular parameters; Fig. S5a, b for organ length and width and Fig. 5g, h for a computation of fold-changes). We, therefore, conclude that it is predominantly a cellular response that controls hypocotyl growth under our conflict-of-interest scenario (Fig. 7a). To explore whether general BR-related cell elongation defects led to the confused phenotypes of some BR pathway mutants, we analysed *bin2-1* mutants, which were among the most severely impaired hypocotyl response to water stress in the dark (Fig. S8a). The data show a most striking impact of *bin2-1* on growth anisotropy, assessed in 2D as length/width (Fig. 5f). Indeed, in a comparison between dark and dark with water stress (darkW), the anisotropy of hypocotyl cells decreased considerably in the wild type (Fig. 5c), but showed no adjustment in *bin2-1* (Fig. 5f). Cell length alone showed the elongation defect typical of *bin2-1* mutants, with a much greater deviation from the wild type under darkW than under dark or light conditions; nonetheless, there was a significant length adjustment to water stress in the dark, even in *bin2-1* (Fig. 5e). These observations suggest that the impaired *bin2-1* hypocotyl response can be attributed to an inability to differentially regulate cell anisotropy in response to the simultaneous withdrawal of light and water.

**Figure 5.**
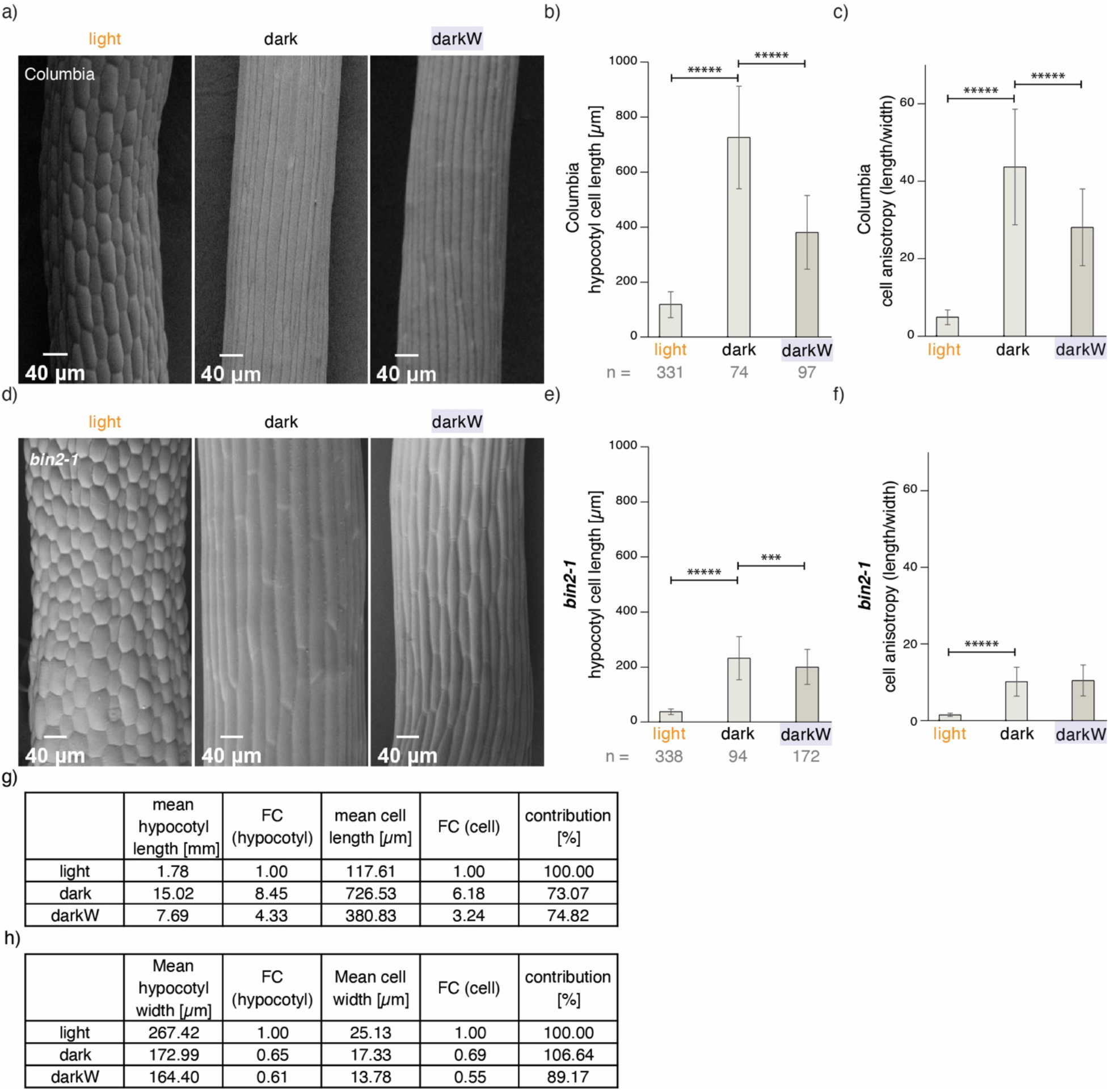
Hypocotyl properties under single versus multiple stress conditions in wild-type and *bin2-1* mutants. Seed were germinated on 1/2MS medium and incubated for ten days under light, dark or darkW conditions. (a-c) Col-0 wild type; (d-f) *bin2-1* semi-dominant allele. Scanning electron micrographs (a, d) were used to assess cellular parameters. (b, e) hypocotyl cell length. (c, f) hypocotyl cell anisotropy measured in 2D as length/width. Notice the highly significant decrease in anisotropy in the wild type (c) but the lack of response in *bin2-1* (f) for darkW as compared to dark. (g, h) A comparison of organ versus cell parameters for the Col-0 wild type under the different environmental conditions, with a computation of fold-changes (FC) with respect to the light condition. Mean cell width significantly (P < 0.00001; not shown) decreased between light and dark and again between dark and darkW (h). Note that, as depicted in the last columns labelled “contributions”, the cellular parameters can account for 73-74% of organ length and 89-106% of organ width. The sample size (n) is given as the number of cells/ number of seedlings that were analysed. P-values were computed with a two-tailed student’s T-test and are represented as follows: ***: 0.001 – 0.0001; *****: < 0.00001. See related Fig. 7.

### BR signalling differentially regulates the timing and extent of cell elongation in the root apical meristem in response to additive stress

It is generally accepted that root growth correlates with the size of the root apical meristem (RAM; Beemster and Baskin 1998). Meristem size was assessed by computing the number of isodiametric and transition cells (González-García et al., 2011; Verbelen et al., 2006; Method S8). In addition, we applied a Gaussian mixed model of cell length to distinguish between short meristematic cells and longer cells in the elongation zone (Fig. S14; Fridman et al., 2021). Meristem size was shortest under water deficit in the dark (Fig. 6a; Fig. S15a,b) and, surprisingly, did not correlate well with final organ length (Fig. 1c; Fig. 6g). We, therefore, measured mature cell length. This was highest in the dark, the condition with the shortest roots (Fig. 6b). Thus, neither meristem size nor mature cell length account for the fold-change in final organ length (Fig. 6g). To address this counterintuitive result, we deployed the CycB1,1: GUS (DiDonato et al., 2004; Method S9) marker, expressed exclusively in cells undergoing M phase (Fig. S15c, S15d). We observed an eight-fold higher number of mitotic cells in darkW as compared to dark conditions (Fig. S15c). This difference was enhanced at an earlier time point (Fig. 6c; Fig. S15c) and could explain the longer organ length but not the smaller meristem size of roots under darkW versus dark (Fig. 6g).

**Figure 6.**
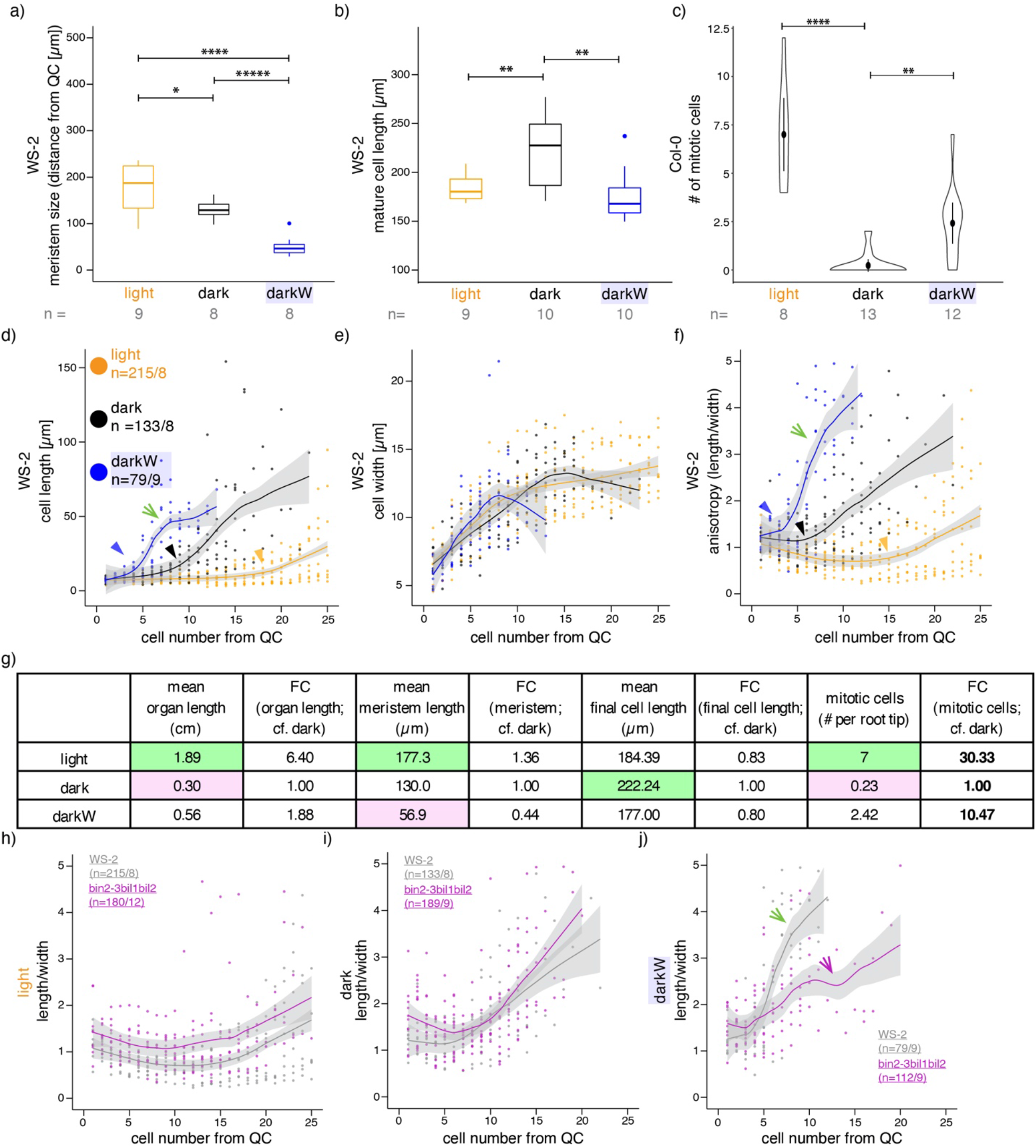
Root meristem properties under single versus multiple stress conditions. Seedlings were grown in the light (orange), dark (black) or dark with −0.4MPa water stress (blue). (a-f) Wild type, with ecotype specified in the panel. (a) Meristem size determined via mixed Gaussian models, as described (Fridman et al., 2021; Fig. S14). (b) Mature cell length, based on the ten most elongated cells for each condition; similar conclusions were reached when the 50 longest cells were used. (c) mitotic index at day 7 based on CycB1,1:GUS (Fig. S15c, S15d; DiDonato et al., 2004). (d-f & h-j) 10 days after incubation, single epidermal cell files were measured, starting at the epidermal/ lateral root cap initials. The fitted lines were generated with Local Polynomial Regression Fitting with the ‘loess’ method in R; grey shading designates the 95 percent confidence interval. (d-f) cell lengths (d), width (e) and anisotropy (in 2D as length/width; f) of consecutive cells as a function of cell number from the quiescent centre (QC); the green arrows point to the steep slope for length (d) and anisotropy (f) under the darkW condition and the arrowheads to the kinks in the curves – the initiation of elongation – under all three conditions. (g) A tabulation of fold-changes (FCs) of measured parameters between different environmental conditions in the wild type (Ws-2, Col-0), with the smallest number highlighted in pink and the largest in green; note that the only FCs that go in same direction as root organ length are for the mitotic index (bold; data depicted in panel c). (h-j) A direct comparison of cell anisotropy under different environmental conditions between the *bin2-3bil1bil2* triple mutant (purple) and the corresponding Ws-2 wild type (grey); notice that the mutant most markedly deviates from the wild type (compare purple versus green arrows in j) in the darkW condition (j), where the steep slope characteristic of the wild type is replaced by a flatter, undulating curve. The sample size (n) is given as the number of seedlings in panels a-c and as the number of cells/ number of seedlings that were analysed in panels h-j. P-values were computed with a two-tailed student’s T-test and are represented as follows: *: 0.05 – 0.01; **: 0.01 – 0.001; ****: 0.0001 – 0.00001, *****: < 0.00001. See related Fig. 7, Fig. S14, Fig. S15 and Fig. S16.

We subsequently looked at cell length, width and anisotropy (computed in 2D as length/width) along single epidermal cell files as a function of distance from the quiescent center. Cell elongation occurred in cells closer to the QC under darkW (blue arrowhead in Fig. 6d) as compared to dark or light conditions (black and orange arrowheads in Fig. 6d). Furthermore, the slopes of both the length and anisotropy curves were steepest for darkW (green arrows in Fig. 6d, 6f). This was also apparent when we looked at the length of the first elongated cell, which was highest in darkW (Fig. S15e). Cell elongation in cells close to the QC translates into an early exit from the root meristem. We conclude that root growth under water deficit in the dark is due not only to increased cell division but also to an early exit from the meristem (Fig. 7b).

**Figure 7.**
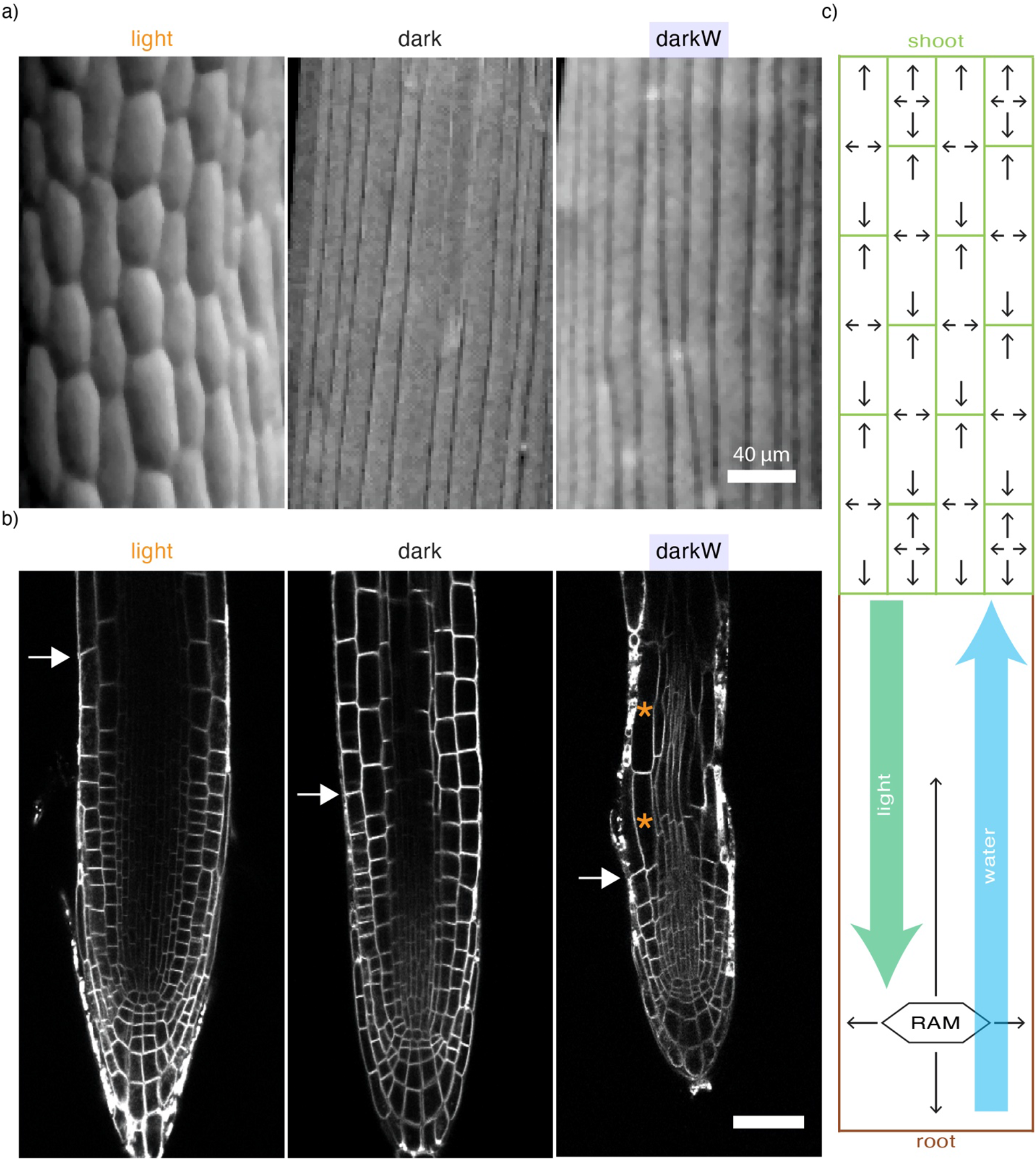
Growth regulation and information flow in response to the availability of water and light. (a) Scanning electron micrographs of wild-type (Col-0) hypocotyls under different environmental conditions. (b) Confocal micrographs of root tips under different environmental conditions (see method S8); white arrows mark the junction between the meristematic and elongation zones (depicted as mean meristem size in Fig. 6a) as assessed by mixed Gaussian model analysis (Fig. S14; Fridman et al., 2021) and orange asterisks mark the highly elongated cells at the very beginning of the elongation zone under darkW. Wildtype (Col-0) for light and dark; P_BRI1_::BRI1-GFP for darkW; FM4-64 signal in all cases. Scalebar: 50 μm. (c) Information-flow diagram as a translation of our empirical findings. The green rectangles in the upper part of the diagram represent hypocotyl cells, and the arrows within each cell depict growth as a decentralized cellular response. The lower part of the diagram (brown-beige) represents the root system, with growth responses (arrows) driven in part by the root apical meristem (RAM). The green arrow depicts a hypothetical hypocotyl to root (basipetal) signal that coordinates trade-offs in organ growth in response to light; this does not take into account the possibility that photoreceptors in the root also perceive and respond to light (Mo et al., 2015). Conversely, the blue arrow depicts a hypothetical root to hypocotyl (acropetal) signal that coordinates trade-offs in organ growth in response to water stress.

We then investigated root meristem properties in *bin2-3bil1bil2,* which had the most aberrant root response to water stress in the dark (Fig. 3e; Fig. S8b). Meristem size and mature cell length followed the same trends in a comparison between *bin2-3bil1bil2* (Fig. S16a, S16b) and the wild type (Fig. 6a, 6b), but the extent of elongation in cells proximal to the QC differed (Fig. S16c). Indeed, *bin2-3bil1bil2* length and anisotropy curves lacked the steep slopes characteristic for darkW in the wild type (compare the green arrows in Fig. 6d, 6f & 6j to the purple arrows in Fig. 6j & Fig. S16c). We conclude that *bin2-3bil1bil2* mutants fail to adjust their root length due to an inability to differentially regulate the elongation of meristematic cells in the root in response to water stress in the dark.

Our observations suggest that root growth under our conflict-of-interest scenario requires root apical meristem function as well as the differential regulation of cell length in the elongation zone. To address this hypothesis, we turned to PLETHORA (PLT) AP2-domain transcription factors, which play a pivotal role in the read-out of the auxin gradient in root tips (Galinha et al., 2007). Indeed, high levels of PLT activity at the stem cell niche promote stem cell identity and maintenance; intermediate levels at the transition zone promote mitotic activity; and low levels in the elongation zone are required for cell differentiation (Galinha et al., 2007). Interestingly, *plt1plt2* mutants (Aida et al., 2004) had an unimpaired hypocotyl response but failed to elongate their roots in response to water stress in the dark (Fig. S15f). Taken together, the cell length and anisotropy curves (Fig. 6) and genetic analyses (Fig. 6; Fig. S15f; Fig. S16) suggest that root length under our different environmental conditions is regulated by (i) the mitotic index, (ii) the timing and extent of cell elongation – translating into the timing of meristematic exit – and (iii) cell geometry. We also conclude that these are differentially modulated to account for increased root length under different environmental conditions (Fig. 6c-e). In addition, an analysis of root meristem properties in *bin2-3bil1bil2* shows that BR signalling is required for the Arabidopsis seedling’s ability to deploy different root growth strategies in response to abiotic stress cues under limiting conditions.

## DISCUSSION

This study establishes screen conditions that enable us to monitor trade-offs between hypocotyl versus root growth in the Arabidopsis seedling. Our screen is based on a conflict-of-interest hypocotyl-versus-root scenario consisting of the simultaneous withdrawal of light and water. A question we addressed was how nutrients, light and water are prioritized in the germinating seedling. We observed a clear priority for hypocotyl growth in search of light over primary root growth in search of nutrients. In contrast to light and nutrients, light and water stress appeared to be equally prioritized in the germinating seedling. Shoot and root lengths were exquisitely fine-tuned to the wavelength and intensity of the light source, and to the severity of water stress. An assessment of organ and cell length suggested that hypocotyl elongation occurred predominantly via cellular elongation. In contrast, root growth appeared to be regulated by a combination of cell division and the timing of exit from the meristem. We have shown that the BR pathway is implicated in hypocotyl versus root trade-offs. BR signalling mutants were most severely impaired in their ability to adjust cell geometry in the hypocotyl and cell elongation as a function of distance from the quiescent centre in the root tips.

It is generally accepted that root length correlates with meristem size. Using kinematic methods, it has also been shown that accelerating root elongation is driven predominantly by an increased number of dividing cells (Beemster and Baskin 1998). Conversely, root growth cessation in response to salt stress has been shown to be a result of decreased cell division, correlating with a reduced meristem size, as well as a reduced mature cell length (West et al., 2004). Therefore, it appears counterintuitive that meristem size and organ length do not correlate in our conflict-of-interest scenario. Questions arise as to why the meristem is smaller under water deficit in the dark even though the mitotic index is higher than in the dark, and how growth is promoted under our additive stress scenarios. An important difference between our conditions and those described by others is that we germinated seed under limiting conditions in the dark in the absence of a carbon source; related studies (such as van der Weele et al., 2000) were carried out in the light or added sucrose to the growth medium in the dark, such that seedlings were not limited with respect to available energy. In this study, we observed growth arrest in the dark, as seen by the low number of mitotic cells in root tips. When water stress was applied in the dark, the mitotic index increased, but the newly produced meristematic cells immediately elongated, thereby exiting the meristem. As a consequence, meristem size remained small despite the increased number of mitotic cells. It appears that what our study shows is a novel paradigm for root growth under limiting conditions, which depends not only on shoot-versus-root trade-offs in the allocation of limited resources, but also on an ability to deploy different strategies for growth in response to abiotic stress cues.

The simplest conceptual framework to explain our observations evokes not only differential regulation of hypocotyl versus root growth, but also hypocotyl to root and root to hypocotyl signalling (Fig. 7c). As water stress was applied exclusively to the root but also impacted hypocotyl growth (Fig. 1e, Fig. S3d), we evoke a root to hypocotyl (acropetal) signal to coordinate trade-offs in organ growth in response to water stress (Fig. 7c blue arrow). Conversely, we postulate that a hypocotyl to root (basipetal) signal coordinates trade-offs in organ growth in response to light (Fig. 7c green arrow). However, and even though photoreceptors are considerably more abundant in the hypocotyl than in the root (van Gelderen et al., 2018), it needs to be borne in mind that photoreceptors in the root could be playing a role in root responses to light or to darkness (Mo et al., 2015). The impact of light was seen under all different wavelengths, suggesting that the light signal acts downstream of all photoreceptors.

There is a fair amount of controversy in the literature regarding the role of BR in cell elongation versus cell division (Nolan et al., 2020). It has, on the one hand, been argued that BR’s impact on cell division (González-García et al., 2011) is a secondary consequence of a primary defect in cell elongation, as cells need to achieve a certain size prior to the initiation of the cell cycle (Jones et al., 2017; Nolan et al., 2020). On the other hand, brassinosteroids have been shown to control QC identity and to impact formative divisions in the root apical meristem (González-García et al., 2011; Kang et al., 2017). Furthermore, high concentrations of BL have been shown to promote cell elongation and an early exit from the root apical meristem (González-García et al., 2011; Chaiwanon and Wang 2015). More recent models for BR signalling in the root show or simulate the greatest impacts on cell anisotropy, growth rates and division plane orientation (Fridman et al., 2021; Graeff et al., 2021). In this study, we addressed the role of BR signalling in cell elongation in the hypocotyl and root tip within the context of growth trade-offs and conflict-of-interest scenarios. While an analysis of cell anisotropy in 3D was not within the scope of this study, our 2D analysis clearly showed that BR signalling was required for an adequate regulation of cell anisotropy in the hypocotyl in response to the simultaneous withdrawal of light and water. In the root, BR signalling mutants were most severely impacted in their ability to differentially adjust cell length as a function of distance from the QC in response to environmental cues.

The BR pathway intersects with light and ABA signalling at many levels. Light is known to inhibit BR biosynthesis, to stabilize or activate BIN2 and to impact BR-responsive transcriptional regulation (Oh et al., 2014; Kim et al., 2011; Li et al., 2020; Li and He 2016; He et al., 2019; reviewed by Nolan et al., 2020; Zhang et al., 2021; Zhao et al., 2022). BIN2 modulates light responses by phosphorylating PIFs, transcriptional regulators inhibited by phytochromes, thereby targeting them for degradation (Bernardo-García et al., 2014; Ling et al., 2017). In addition, the interaction between BIN2 and its substrates is regulated by photoreceptors or light signalling components (Ling et al., 2017; He et al., 2019). The BR pathway has also been implicated in drought responses and ABA signalling (Cui et al., 2019; Nolan et al., 2017). BIN2 is a target of PP2C phosphatases in the abscisic acid pathway and mediates drought responses by targeting ABA signalling components and drought or desiccation responsive transcription factors (Cai et al., 2014; Hu and Yu 2014; Jiang et al., 2019; Wang et al., 2014; Wang et al., 2018; Xie et al., 2019). Furthermore, BIN2 homologues have been implicated in root responses to osmotic stress (Dong et al., 2020). In addition to its role in light and drought responses, the BR pathway has been implicated in the growth versus defence trade-off (Lozano-Durán and Zipfel 2015; Ortiz-Morea et al., 2020) and in regulating hypocotyl elongation in response to far-red light and salt stress (Hayes et al., 2019). Studies on responses to abiotic stress factors have typically addressed growth arrest or trade-offs between growth and acclimation (Bechtold and Field 2018). Indeed, root growth is inhibited by, for example, phosphate deprivation or salt stress (Balzergue et al., 2017; West et al., 2004). Recent efforts have addressed strategies for engineering drought resistant or tolerant plants that do not negatively impact growth (Fàbregas et al., 2018; Yang et al., 2019). In contrast to other studies, here we look at two abiotic stress factors that promote organ growth. Indeed, hypocotyl growth is promoted by darkness or low light and primary root growth by water deficit in this study.

In the judgement and decision-making model for plant behaviour put forth by Karban and Orrock (2018), signal integration might be considered integral to judgement. In decision theory one could refer to signal integration as an assessment of the “state of the world”, although in our case this state is integrated with endogenous developmental signals as well. In our conflict-of-interest scenario, the “confused” phenotypes of BR signalling mutants are indicative of an integral role in decision-making *per se.* Whether judgement and decision making can be distinguished from each other empirically remains unclear. As BR signalling regulates cell anisotropy and the timing and extent of cell elongation in the hypocotyl and root apical meristem, it may play a role not only in signal integration but also in the execution of decisions (or in an implementation of the action). Thus, this study does not enable us to empirically distinguish between decision making on the one hand and signalling and execution on the other.

Future experiments will address where in the seedling decision-making processes occur, the nature, interdependence and movement of the acro- and basipetal signals, and how these facilitate shoot to root communication to fine tune trade-offs between root versus shoot growth. 3D imaging will be required to assess the impact of abiotic stress and/or of BR signalling on different cell files or tissue layers in the root (see Hacham et al., 2011; Fridman et al., 2014; Fridman et al., 2021; Graeff et al., 2021). Similarly, time-lapse imaging will be required for temporal resolution. As a limited budget is an essential component of our screen conditions, the role of energy sensing and signalling (Baena-González and Hanson 2017) in growth trade-offs will need to be elucidated. In addition, phosphoproteomics may enable us to better understand BIN2 targets under our conflict-of-interest-scenario. The screen has broader implications for plant responses to multiple stress parameters applied simultaneously. Indeed, with changing climate and mounting degrees of uncertainty, a possibly comforting outcome of our screen is that although we actively looked for mutants with “confused” phenotypes, we were, with the exception of selected BR signalling mutants, hard put to uncover any. Rather, our overall conclusion is that even mutants with very severe growth defects were, when faced with extreme multiple stress conditions, by and large capable of adjusting their shoot to root growth trade-offs to optimize their chances of survival upon germination.

## Supporting information

Kalbfuss et al., 2022, supplemental information

## ACKNOWLEDGEMENTS

We thank Prof. Grill for his support. Alexander Christmann, Mikhal Kepka and other members of the Botany department provided EMS-mutagenized seed and made useful suggestions. Caroline Klaus tended to our plants in the green house and made valuable phenotypical observations. We are exceedingly grateful to Andreas Czempiel and Knut Thiele for technical assistance. Fiona Pachl initiated the positional cloning of B1. We thank Leon Assaad for insights into decision theory. Laura Gräbener, Manuel Jeller and Tobias Weiser contributed to this study as undergraduate students. Urs Schmidhalter provided expertise for nutrient stress. We thank Frej Tulin for a critical appraisal of the manuscript. The NASC stock center distributed public seed stocks and Zhiyong Wang, Joanne Chory, Jorge José Casal and John Celenza shared published resources. We thank Roman Meier at the TUMmesa facility, directed by Leonardo Teixera, for supporting us with optimal growth conditions for our plants. We gratefully acknowledge the WZW/TUM Centre for Advanced Light Microscopy (CALM), headed by Ramon Torres-Ruiz, for unlimited access to confocal microscopes. The authors declare no conflict of interest.

## AUTHOR CONTRIBUTIONS

N.K. designed, performed and supervised mutant screening experiments, analyzed, quantified and interpreted data, assembled all figure panels and assisted in drafting sections of the manuscript.

A. S. performed, analysed and interpreted CSLM analysis of root meristem parameters.

M.K. designed, performed, analysed and interpreted light and GUS-staining experiments.

L.L. studied the impact of a gradient of water stress on wild-type ecotypes.

F.G.H. studied the BR pathway and analyzed, quantified and interpreted CSLM data on the RAM.

B. B., S.P. and S.S analysed SEM datasets as well as water stress responses in the light.

K.S. performed nutrient stress experiments.

K.S. and L.L. positionally cloned the B1 mutant.

C. W., C.F., C.S. and M.T. performed, analysed and interpreted screening experiments

K.F. performed mannitol and salt stress experiments and helped establish the method.

J.H. helped with literature searches and drafted a section of the discussion

E. F. performed and supervised SEM image acquisition.

W.P. contributed to the analysis of cell growth dynamics and to the information flow diagram.

C.B. designed, supervised, analysed and interpreted light and qPCR experiments.

F. F.A. acquired funding, designed the screen, performed and analysed experiments. (co)supervised all experiments, wrote and revised the manuscript with input from all co-authors.

All data that support the conclusions of this study are available upon request. The author responsible for distribution of materials in accordance with the policy described in the Instructions for Authors is: Farhah Assaad.

## SUPPORTING INFORMATION

**Fig. S1** The impact of nutrient stress (-K, -N, -P) on hypocotyl versus root growth in dark-grown seedlings.

**Fig. S2** Hypocotyl versus root growth in response to osmotic stress and salt stress in dark-grown seedlings.

**Fig. S3** Tradeoffs between hypocotyl and root growth in response to water stress.

**Fig. S4** Impact of different light conditions and light signaling on hypocotyl versus root growth.

**Fig. S5** Impact of water stress on hypocotyl length, width and volume in dark-grown seedlings.

**Fig. S6** Light responses in the B1 mutant and comparison to *bin2-1.*

**Fig. S7** The response of BR pathway mutants to water stress in the dark: violin plots

**Fig. S8** The response of BR pathway mutants to water stress in the dark: response quotients and volcano plots.

**Fig. S9** Seed weights in selected BR mutants.

**Fig. S10** Light responses in BR pathway mutants: bar graphs.

**Fig. S11** Light responses in BR pathway mutants: response quotients and volcano plots.

**Fig. S12** Expression of light responsive gene *LHCB1.2*

**Fig. S13** Responses of selected BR mutants to water stress in the light

**Fig. S14** Root apical meristem size under different environmental conditions

**Fig. S15** Root apical meristem properties under different environmental conditions

**Fig. S16**. Root meristem properties in *bin2-3bil1bil2* under multiple stress conditions.

**Table S1** Lines used in this study

**Table S2** Segregation analysis of the B1 mapping population

**Method S1** Composition of nutrient stress plates

**Method S2** Supplemental information on light experiments

**Method S3** Preparation of PEG plates

**Method S4** Seed handling for screen

**Method S5** Picking, scanning, phenotyping and genotyping

**Method S6** Positional Cloning

**Method S7** Real-time PCR analysis

**Method S8** Confocal microscopy and root apical meristem properties

**Method S9** GUS staining

