## Supplementary material for "A role for brassinosteroid signaling in decision-making processes in the Arabidopsis seedling": Kalbfuss et al., 2022, supplemental information

Article title: A role for brassinosteroids in decision-making processes in the Arabidopsis seedling

**SUPPLEMENTAL INFORMATION**

**Fig. S1** The impact of nutrient stress (-K, -N, -P) on hypocotyl versus root growth in dark-grown seedlings.
**Fig. S2** Hypocotyl versus root growth in response to osmotic stress and salt stress in dark-grown seedlings.
**Fig. S3** Tradeoffs between hypocotyl and root growth in response to water stress**.**
**Fig. S4** Impact of different light conditions and light signaling on hypocotyl versus root growth.
**Fig. S5** Impact of water stress on hypocotyl length, width and volume in dark-grown seedlings.
**Fig. S6** Light responses in the B1 mutant and comparison to *bin2-1*.
**Fig. S7** The response of BR pathway mutants to water stress in the dark: violin plots
**Fig. S8** The response of BR pathway mutants to water stress in the dark: response quotients and volcano plots.
**Fig. S9** Seed weights in selected BR mutants.
**Fig. S10** Light responses in BR pathway mutants: bar graphs**.**
**Fig. S11** Light responses in BR pathway mutants: response quotients and volcano plots**.**
**Fig. S12** Expression of light responsive gene *LHCB1.2*
**Fig. S13** Responses of selected BR mutants to water stress in the light
**Fig. S14** Root apical meristem size under different environmental conditions
**Fig. S15** Root apical meristem properties under different environmental conditions
**Fig. S16**. Root meristem properties in *bin2-3bil1bil2* under multiple stress conditions.
**Table S1** Lines used in this study
**Table S2** Segregation analysis of the B1 mapping population
**Method S1** Composition of nutrient stress plates
**Method S2** Supplemental information on light experiments
**Method S3** Preparation of PEG plates
**Method S4** Seed handling for screen
**Method S5** Picking, scanning, phenotyping and genotyping
**Method S6** Positional Cloning
**Method S7** Real-time PCR analysis
**Method S8** Confocal microscopy and root apical meristem properties
**Method S9** GUS staining

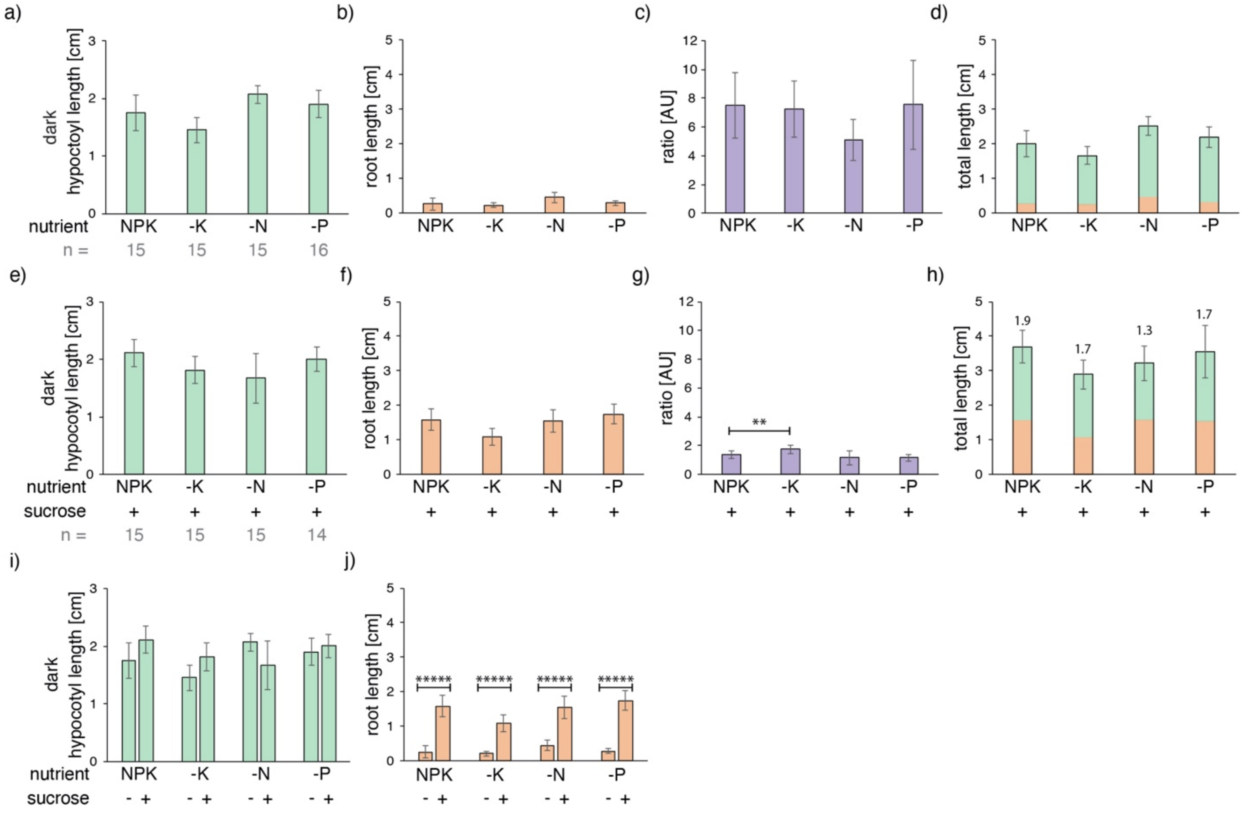


**Fig. S1 The impact of nutrient stress (-K, -N, -P) on hypocotyl versus root growth in dark-grown seedlings.** Col-0 seed were plated on NPK media with or without (-) K, N or P and incubated for seven days in the dark. (a-d) germination in the absence of a carbon source. Panels (a-d) are from the same experiment. Minor fluctuations were detected, but no clear tradeoffs between hypocotyl and root growth were observed when nutrient stress was applied to dark-grown seedlings. Based on the long hypocotyls (a) and short roots (b), there is a clear priority for light (translating into hypocotyl growth) over nutrients (translating into root growth) in the germinating seedling as seen in the hypocotyl/root ratio (c) and the total length (d). (e-h) germination in the presence (+) of 1 % sucrose as a carbon source. The presence of sucrose as a carbon source increased the total length of the seedlings up to two-fold (h); numbers above the columns are fold-changes compared to the same condition without sucrose. (i, j) Panels in (i, j) depict the impact of a carbon source. In the presence (+) of a carbon source, hypocotyl length remained fairly constant but the roots were able to grow longer (up to 6-fold increase in root length, as compared to the absence (-) of sucrose). Thus, there was no clear tradeoff, by which we refer to the growth of one organ at the expense of another. As the genetic screen was designed to mimic limiting conditions, we omitted sucrose from the media in all further experiments. The number (n) of seedlings measured per condition is in grey below the graph. *P*-values were computed with a two-tailed student's *T*-test and are represented as follows: **: 0.01 - 0.001; *****: < 0.00001. Related to Fig. 1.

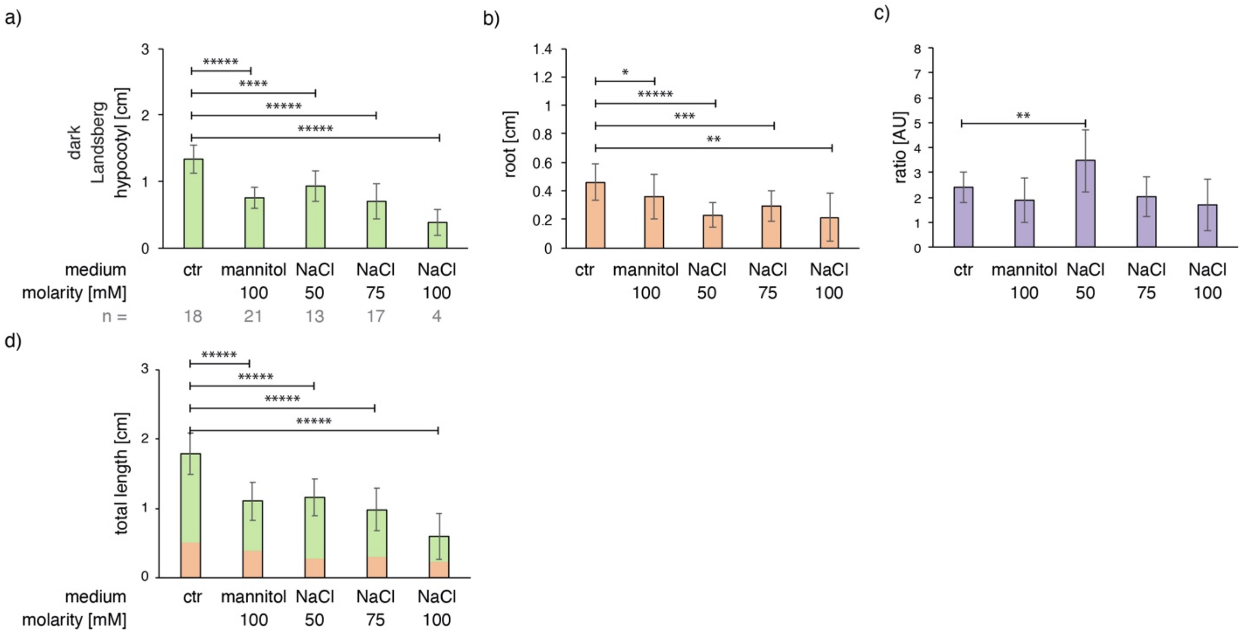


**Fig. S2 Hypocotyl versus root growth in response to osmotic stress and salt stress in dark-grown seedlings.** Hypocotyl/root ratio of seedlings (Ler wild type) germinated under osmotic (100mM mannitol) or salt stress (50-100mM NaCl) in the dark in the absence of a carbon source. No reproducible differences in the ratio were observed (P > 0.5). The number (n) of seedlings measured per condition is in grey below the graph. P-values were computed with a two-tailed student's T-test and are represented as follows: *: 0.05 - 0.01; **: 0.01 - 0.001; ***: 0.001 - 0.0001; ****: 0.0001 - 0.00001, *****: < 0.00001. Related to Fig. 1

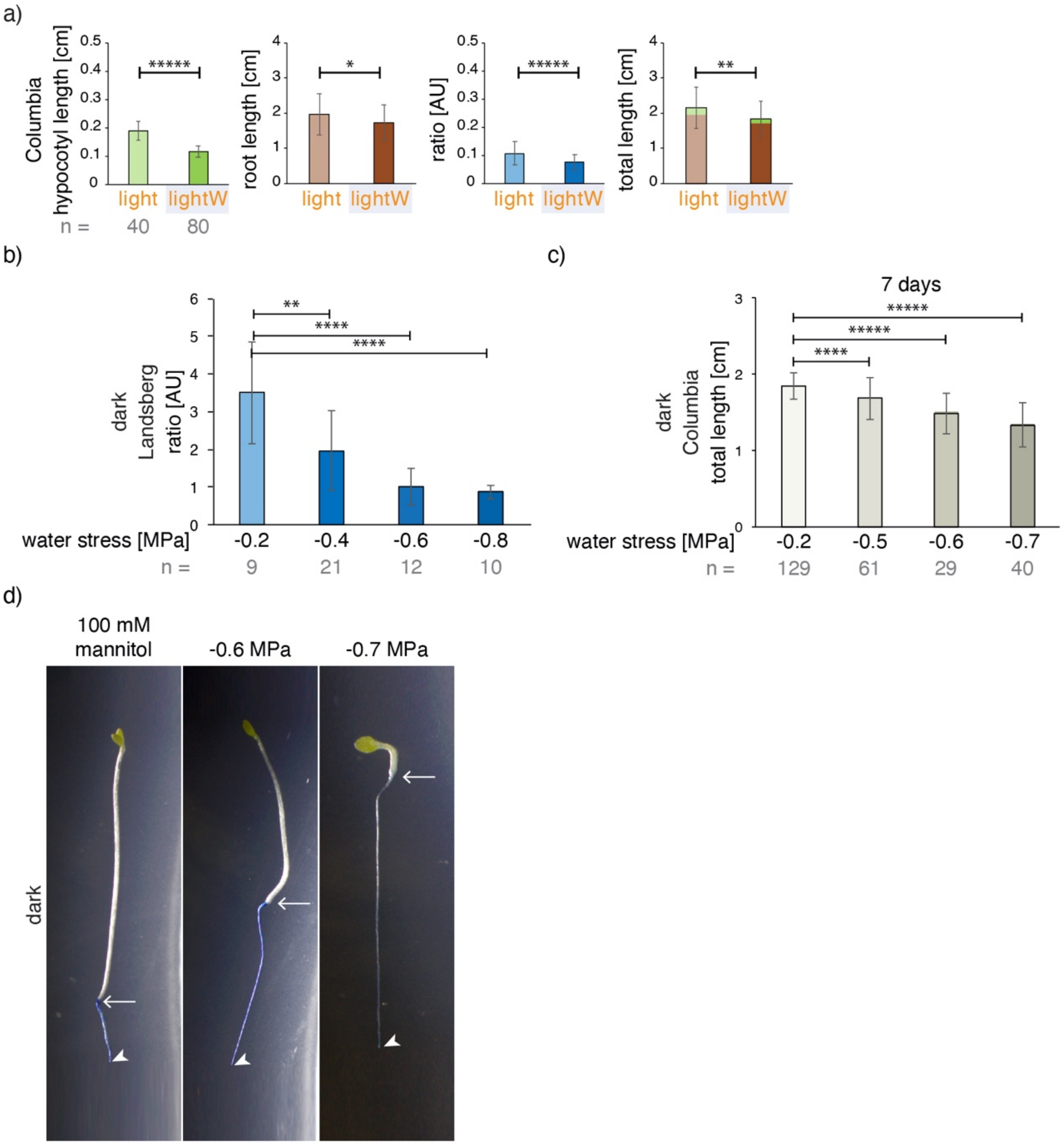


**Fig. S3 Tradeoffs between hypocotyl and root growth in response to water stress.** (a) Hypocotyl/root ratio of seedlings (Col wild type) germinated in the light with (lightW) or without (light) water stress (PEG -0.4MPa); the clearest response to water stress in the light is a reduction in hypocotyl length (left panel). This is consistent with the observations of van der Weele et al. (2000) on the use of PEG in the light. (b) Seed (Ler) were geminated in the dark on MS medium (- 0.2 MPa) with a gradient of water stress ranging from -0.2 to -0.8 MPa. The hypocotyl/root ratio of Ler seedlings is similar to that of Col-0 (Fig. 1d): water stress applied in the dark increases root length at the expense of hypocotyl length, giving rise to a decrease in the hypocotyl/root ratio. (c) The length of wild type (Col-0) seedlings was seen to decrease with increasing water stress after seven (but not after ten days; cf. Fig. 1e), and this decrease was observed in control experiments to be due to delayed germination induced by water stress. (d) Roots were stained with methanol blue; arrows point to the junction between the hypocotyl and root and arrowheads to the end of the root. The number (n) of seedlings measured per condition is in grey below the graph. P-values were computed with a two-tailed student's T-test and are represented as follows: *: 0.05 - 0.01; **: 0.01 - 0.001; ***: 0.001 - 0.0001; ****: 0.0001 - 0.00001, *****: < 0.00001. Related to Fig. 1 and S5.

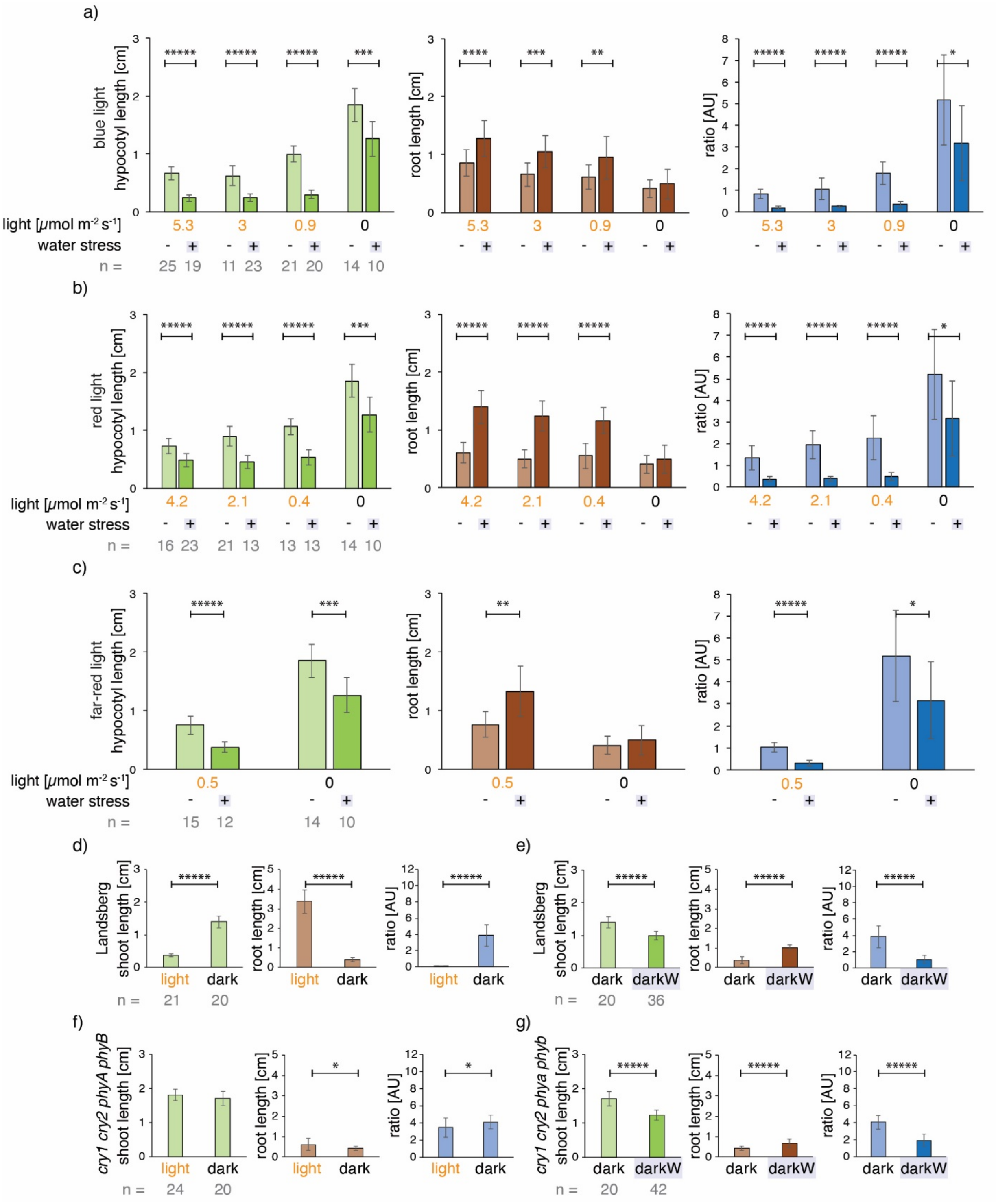


**Fig. S4 Impact of different light conditions and light signaling on hypocotyl versus root growth.** (a-c) Wild-type (Col-0) seedlings were germinated on MS media under different light conditions, with or without water stress. (a) Blue light at varying intensities ranging from 5.3 to 0 μmol m$^{-2}$ s$^{-1}$. (b) Red light at varying intensities ranging from 4.2 to 0 μmol m-2 s-1. (c) Far-red light at 0.5 μmol m-2 s-1, compared to dark grown seedlings. A decreasing intensity gradient of red light, as well as low levels of far-red light, increases hypocotyl length at the expense of root length as seen in the hypocotyl/root ratio. (d-g) Quadruple *cry1 cry2 phyA phyB* mutant seedlings (f,g) had no hypocotyl response to light (f; white light intensity: 250 μmol m$^{-2}$

$s^{-1}$, compared to dark conditions) but had highly significant responses to water stress applied in the dark (g); corresponding Ler wild-type (d,e). The number (n) of seedlings measured per condition is in grey below the graph. Benjamini–Hochberg corrected P-values are represented as follows: *: 0.05 - 0.01; **: 0.01 - 0.001; ***: 0.001 - 0.0001; ****: 0.0001 - 0.00001, *****: < 0.00001. Related to Fig. 1.

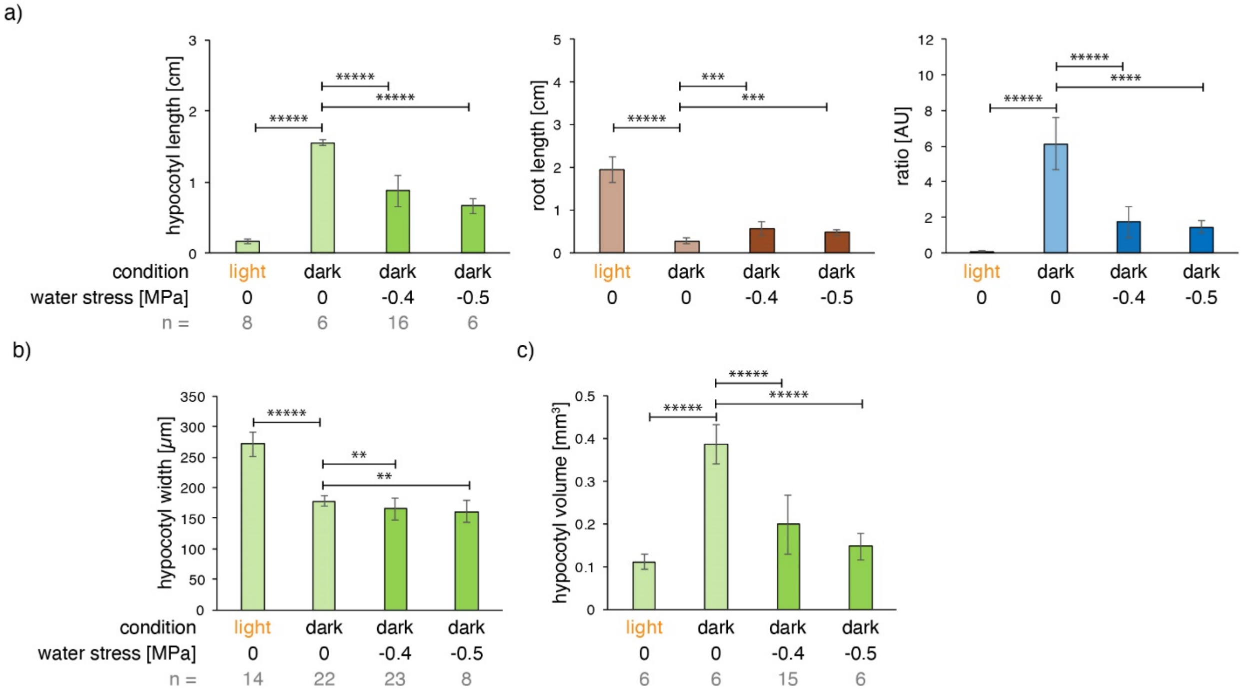


**Fig. S5 Impact of water stress on hypocotyl length, width and volume in dark-grown seedlings**. Wild-type seed (Col-0) were germinated on ½ MS in the dark with or without water stress. (a) Hypocotyl and root lengths, as well as hypocotyl/root ratio. Note that seedlings germinated under water stress in the dark have shorter hypocotyls and longer roots than in the dark, giving rise to a decreased hypocotyl/root ratio. (b) Hypocotyl width; hypocotyls are wider in light than in the dark; water stress in the dark results in a decrease in hypocotyl width. (c) Hypocotyl volume; the volume increases in the dark (cf light) and decreases in response to water stress in the dark. The number (n) of seedlings measured per condition is in grey below mean ±StDev bar graphs. *P*-values were computed with a two-tailed student's *T*-test and are represented as follows: **: 0.01 - 0.001; ***: 0.001 - 0.0001; ****: 0.0001 - 0.00001, *****: < 0.00001. Scale bars = 1mm. See related Fig. 1 and Fig. S3.

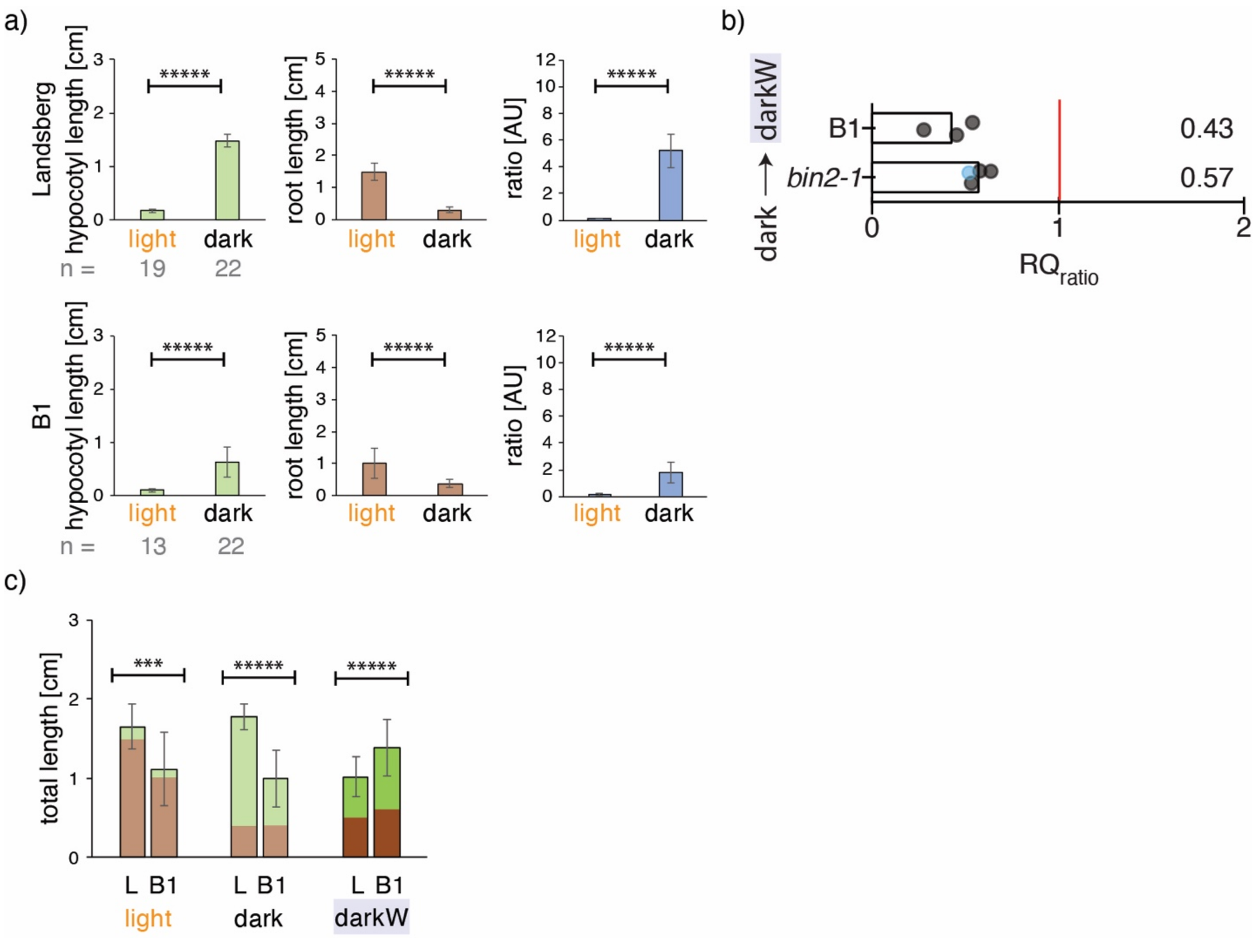


**Fig. S6 Light responses in the B1 mutant and comparison to *bin2-1*.** (a) Light responses were attenuated in B1, with a shorter hypocotyl and longer root in the dark. (b) RQratios (see main methods) were comparable and attenuated in both B1 and *bin2-1*. (c) The total length is depicted as the hypocotyl (top, green) and root (bottom, beige). B1 was shorter than Ler (L) in the light and in the dark, but not under darkW conditions. The number (n) of seedlings measured per condition is in grey below the graph. P-values were computed with a two-tailed student's T-test and are represented as follows: *: 0.05 - 0.01; **: 0.01 - 0.001; ***: 0.001 - 0.0001; ****: 0.0001 - 0.00001, *****: < 0.00001. Related to Fig. 2.

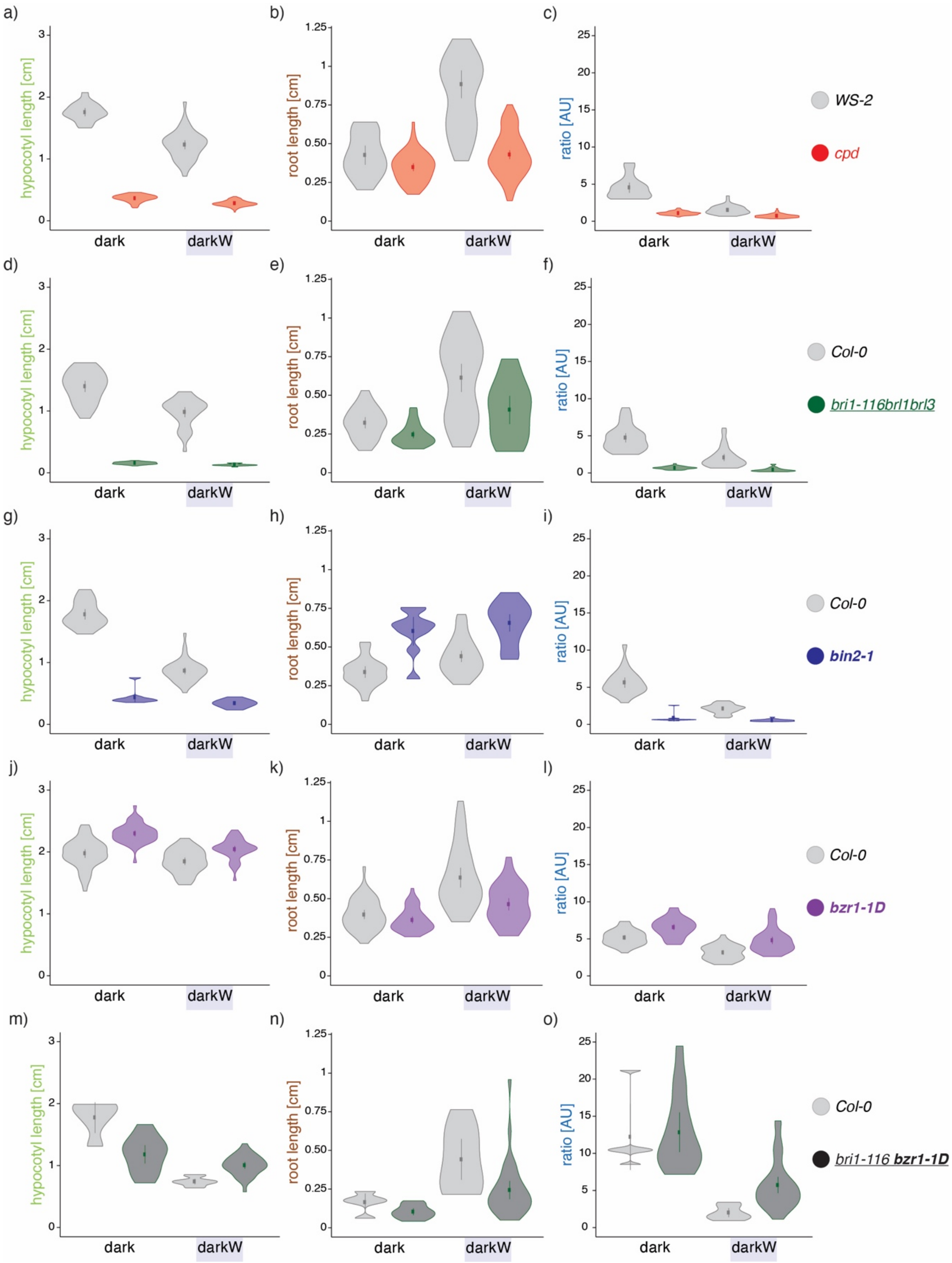


**Fig. S7 The response of BR pathway mutants to water stress in the dark: violin plots.** Violin plots of the hypocotyl, root and ratio responses of the BR pathway mutants shown in Fig. 3, with the corresponding wild-type ecotype as reference. The dot represents the mean and the line the 95% confidence interval. Note the high variance of the wild-type (Col-0) root response under darkW. Mutant alleles are described in Table S1; null alleles are depicted in regular font, semi-dominant or dominant in bold and higher order mutants are underlined. Sample sizes and statistics are as in Fig. 3.

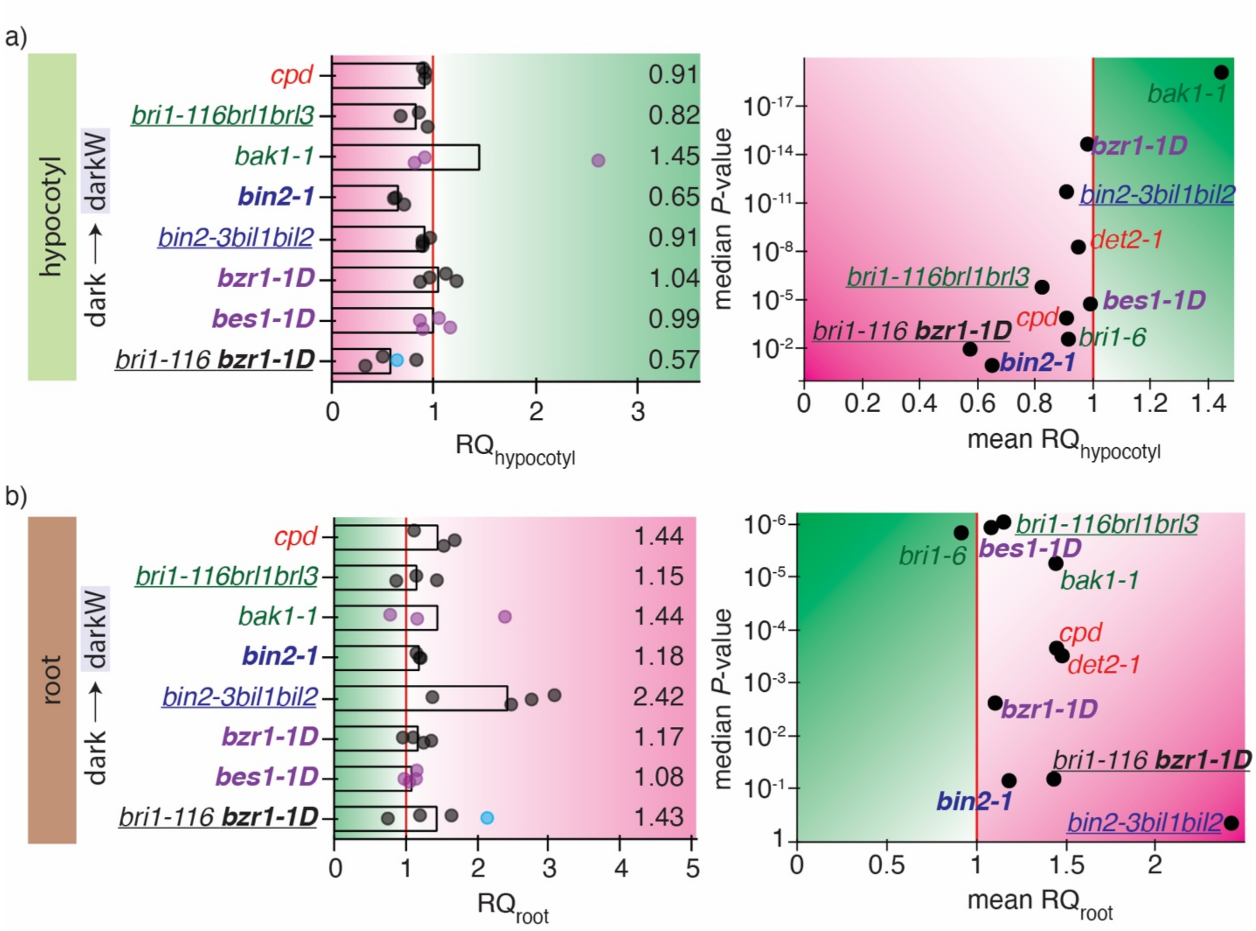


| | RQ dark/darkW | | |
|---|---|---|---|
| | hypocotyl | root | ratio |
| attenuated response | <0.8 | >1.2 | <0.8 |
| normal response | 0.8-1.2 | 0.8-1.2 | 0.8-1.2 |
| exaggerated response | >1.2 | <0.8 | >1.2 |

**Fig. S8 The response of BR pathway mutants to water stress in the dark.** Response quotients (RQ, left) and volcano plots (right) of the hypocotyl and root responses water stress in the dark. RQs are normalized to the wild-type ratio quotient; a value of 1 (vertical red line) indicates that the response to a shift from dark to darkW is similar to that of the respective wild-type ecotype. Each replicate is represented by a dot; purple dots are for initial and grey dots for optimized screen conditions; blue dots are for data from SEM measurements. Volcano plots with the mean RQ depicted on the left on the *X*-axis and the *P*-Value of the response on the *Y*-axis (negative log scale; a median of all replicates was used). Null alleles are depicted in regular font, semi-dominant or dominant in bold and higher order mutants are underlined. (a) Hypocotyl responses to dark versus darkW conditions. Note that *bri1-116 bzr1-1D* mutants have a severely attenuated and *bin2-1* an attenuated hypocotyl response, $RQ_{hypocotyl}$. (b) Root responses to dark versus darkW conditions. Note that the triple *bin2bil1bil2* knock out has the strongest $RQ_{root}$ phenotype. Thresholds used to interpret the results are tabulated at the bottom of the figure; magenta color indicates an attenuated and green an exaggerated response. Related to Figures 3 + 4.

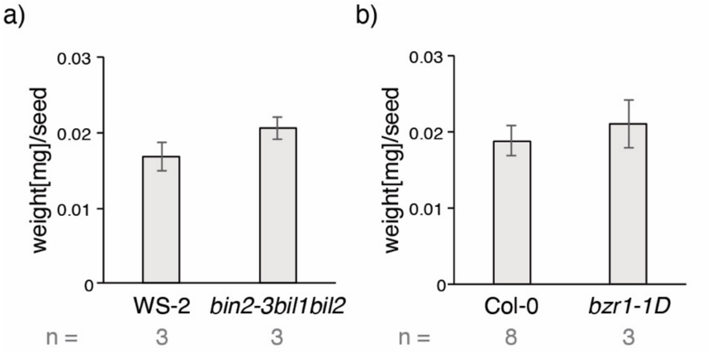


**Fig. S9 Seed weights in selected BR mutants.** The number (n) of seed bags analysed per condition is in grey below mean ±StDev bar graphs. Each seed-bag contained on average 397 seed. *P*-values were computed with a two-tailed student's *T*-test and were non-significant: >0.05. Related to Fig. 5 and 6.

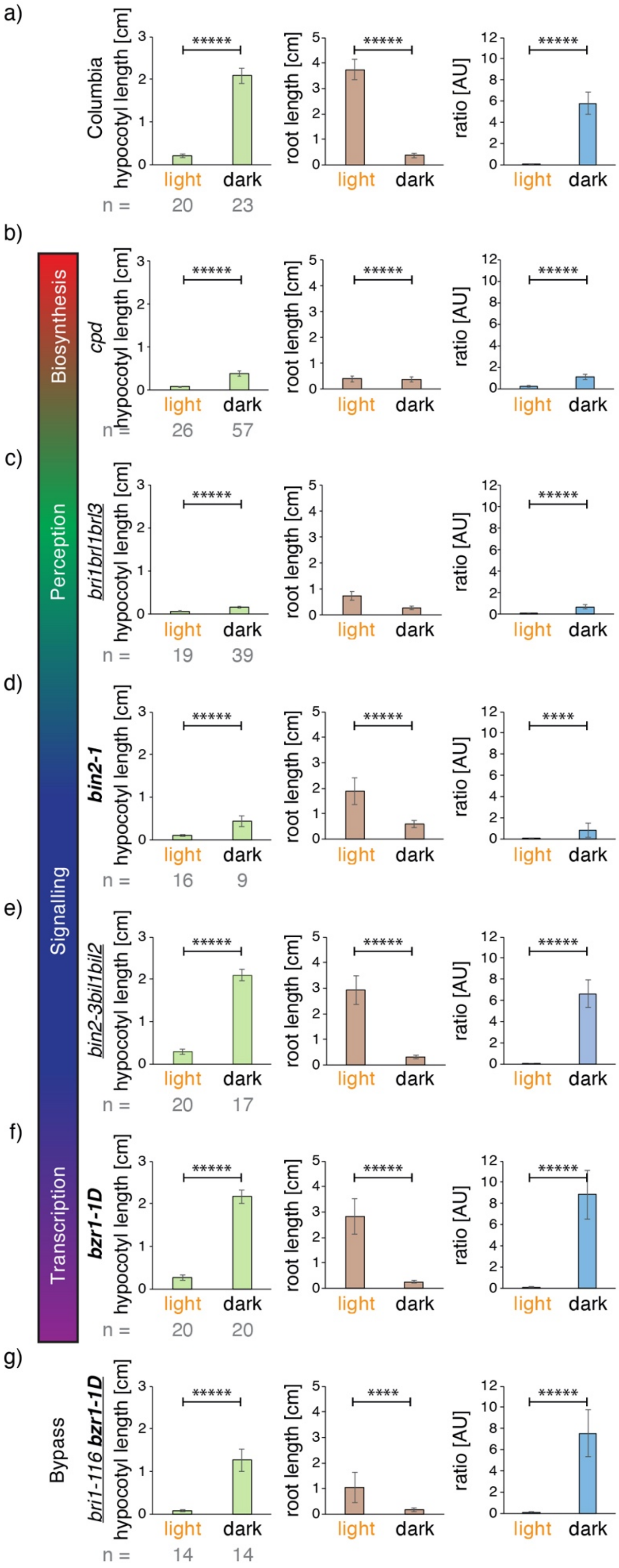


**Fig. S10 Light responses in BR pathway mutants: bar graphs.** Seedlings were germinated on ½ MS in the light and dark (a) Col-0 (wild type). (b) BR biosynthesis mutant *cpd*. (c) BR perception mutant *bri1brl1brl3*(d) BR signaling mutant *bin2-1* (a semidominant gain of function allele). (e) *bin2-3bil1bil2* tripple knockout; (f) Transcription factor mutant *bzr1-1D*, a dominant allele. (g) BIN2 bypass mutant *bri-116 bzr1-1D*. (a-m) Null alleles are depicted in regular font, semi-dominant or dominant in bold and higher order mutants are underlined. Note that all mutants had significant responses to light versus dark conditions. At least 3 experiments were performed for each line, and a representative one is shown here on the basis of RQ and P values (see Fig. S11). The number (n) of seedlings measured per condition is in grey below the mean ±StDev bar graphs. *P*-values were computed with a two-tailed student's *T*-test and are represented as follows: *: 0.05 - 0.01; **: 0.01 - 0.001; ***: 0.001 - 0.0001; ****: 0.0001 - 0.00001, *****: < 0.00001. For mean RQ values and median *P*-values see Fig. S11. Ecotypes are described in Table S1. Related to Figures 3 + 4.

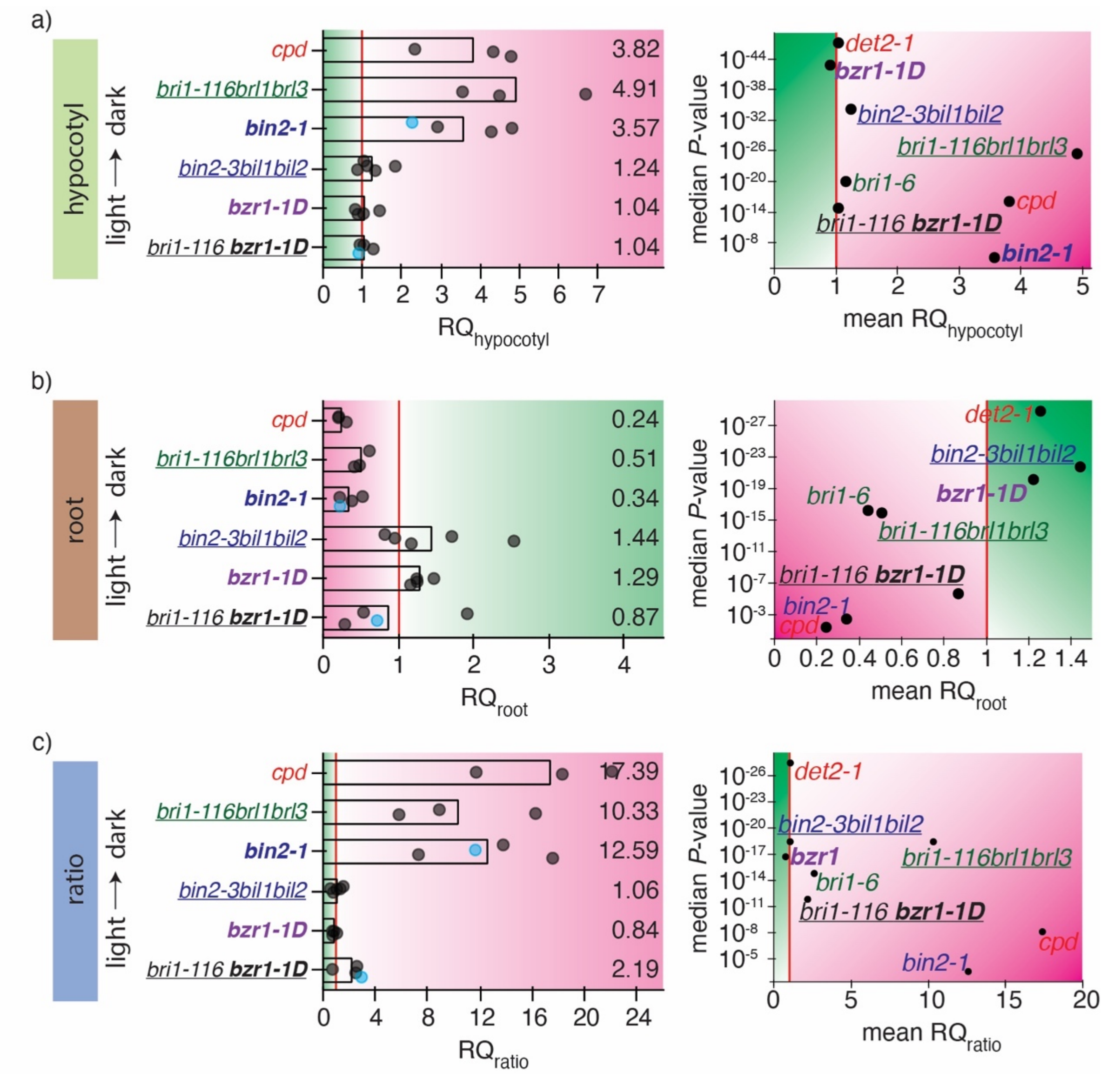


| | RQ light/dark | | |
|---|---|---|---|
| | hypocotyl | root | ratio |
| attenuated response | >1.2 | <0.8 | >1.2 |
| normal response | 0.8-1.2 | 0.8-1.2 | 0.8-1.2 |
| exaggerated response | <0.8 | >1.2 | <0.8 |

**Fig. S11 Light responses in BR pathway mutants. Response quotients** (RQ, left) and **volcano plots** (right) of the hypocotyl, root or ratio responses to light versus dark. RQs are normalized to the wild-type ratio quotient; a value of 1 (vertical red line) indicates that the response to a shift from light to dark is similar to that of the respective wild-type ecotype. Each replicate is represented by a dot; grey dots are for optimized screen conditions; blue dots are for data from SEM measurements. Volcano plots with the mean RQ depicted on the left on the *X*-axis and the *P*-value of the response on the *Y*-axis (negative log scale; a median of all replicates was used). Null alleles are depicted in regular font, semi-dominant or dominant in bold and higher order mutants are underlined. (a) Hypocotyl responses to light versus dark conditions. Note that *cpd, bri1brl1brl3* and *bin2-1* mutants have a severely attenuated hypocotylhypocotyl response $RQ_{hypoctyl}$. (b) Root responses to light versus dark conditions. Note that *cpd,* and *bin2-1* mutants have the strongest $RQ_{root}$ phenotype, and that *bin2-1* gain of function and *bin2-3bil1bil2* loss of function mutants have opposite root phenotypes. (c) Hypocoytl/root ratio responses to light versus dark conditions. In all three volcano plots, *cpd* and *bin2-1* mutants are the most severely impaired (most attenuated response (RQ), lowest P-value). Thresholds used to interpret the results are tabulated at the bottom of the figure; magenta color indicates an attenuated and green an exaggerated response. Related to Figures 3 + 4.

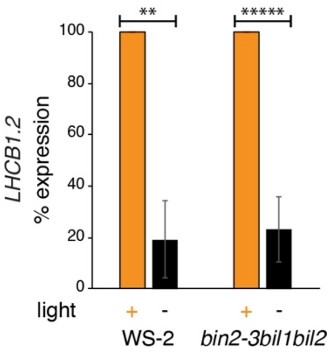


**Fig. S12 Expression of light responsive gene *LHCB1.2*** in Ws-2 wild-type (WT) and *bin2-3bil1bil2* mutant seedlings. Seed were germinated on 1/2 MS plates and incubated in the light (orange; +) or dark (black; -) for 10 days. Transcript abundance was determined by qRT-PCR with *Ubiquitin-protein ligase-like protein* as a reference for normalization (see Methods S7). Expression in the light was set at 100% for both genotypes. Gene expression was significantly downregulated in the dark, in both the wild-type and in *bin2-3bil1bil*2 mutants. Data represent means ± StDEV of three independent experiments, where each measurement was based on three technical replicates. Three biological replicates were used for the mutant and two biological replicates for the wild type. *P*-values were computed with a two-tailed student's *T*-test and are represented as follows: **: 0.01 - 0.001; *****: < 0.00001.

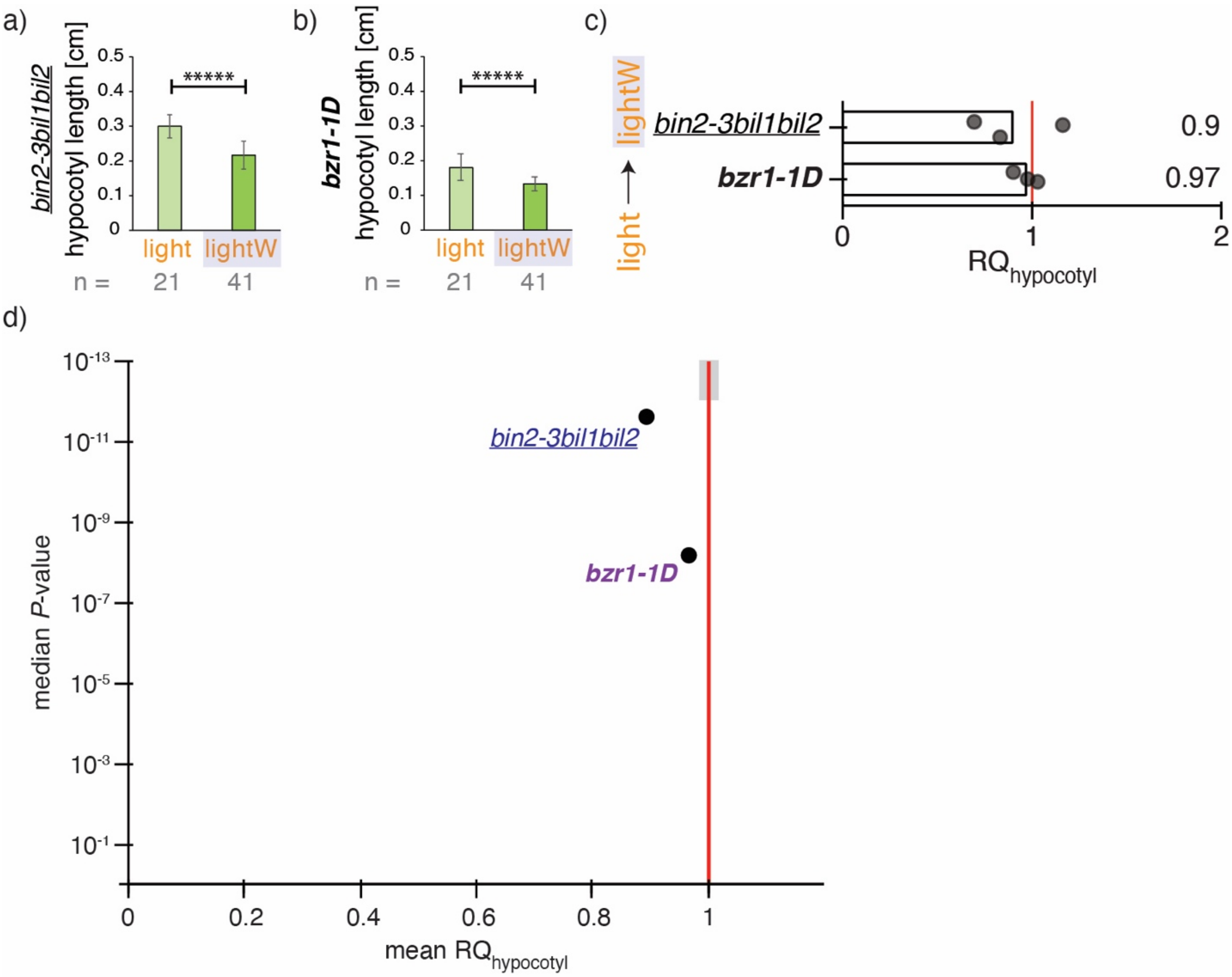


**Fig. S13 Responses of selected BR mutants to water stress in the light** (lightW). Hypocotyl responses of (a) *bin2-3bil1bil2* tripple knockout; (b) Transcription factor mutant *bzr1-1D*, a dominant allele. The number (n) of seedlings measured per condition is in grey below the graph. *P*-values were computed with a two-tailed student's *T*-test and are represented as follows: *****: < 0.00001 (c) $RQ_{hypocotyl}$ response quotient of the hypocotyl under light/lightW conditions, normalized to the wild-type response quotient; a value of 1 (vertical red line) indicates that the response to a shift from light to lightW is similar to that of the respective wild-type ecotype. Each replicate is represented by a dot. (d) Volcano plot with the mean $RQ_{hypocotyl}$ depicted in (c) on the *X*-axis and the median *P*-Value of the response on the *Y*-axis (negative log scale; a median of all replicates was used). Related to Figures 3 + 4.

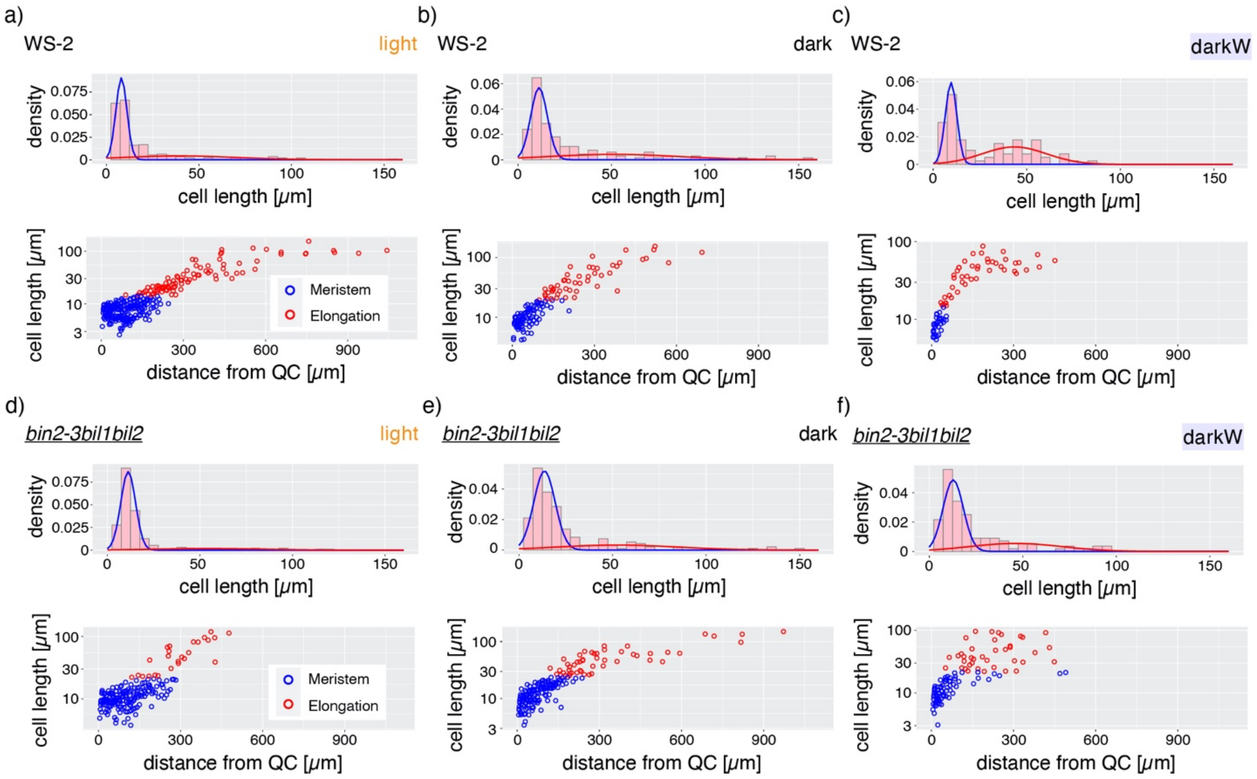


**Fig. S14 Root apical meristem size under different environmental conditions.** As described by Fridman et al., 2021, we used the expectation maximization algorithm as implemented in the mixtools R package to fit a two-Gaussian mixture model to the cell length parameter in each condition. This captures two populations of short versus long cells. Cells with a probability > 0.8 of being in the short-length Gaussian were considered to be meristematic cells (blue). Cells outside the meristem were considered as being elongating cells (red). (a-c) wild-type (Ws-2). (d-f) *bin2-3bil1bil2* triple null BR signaling mutant.

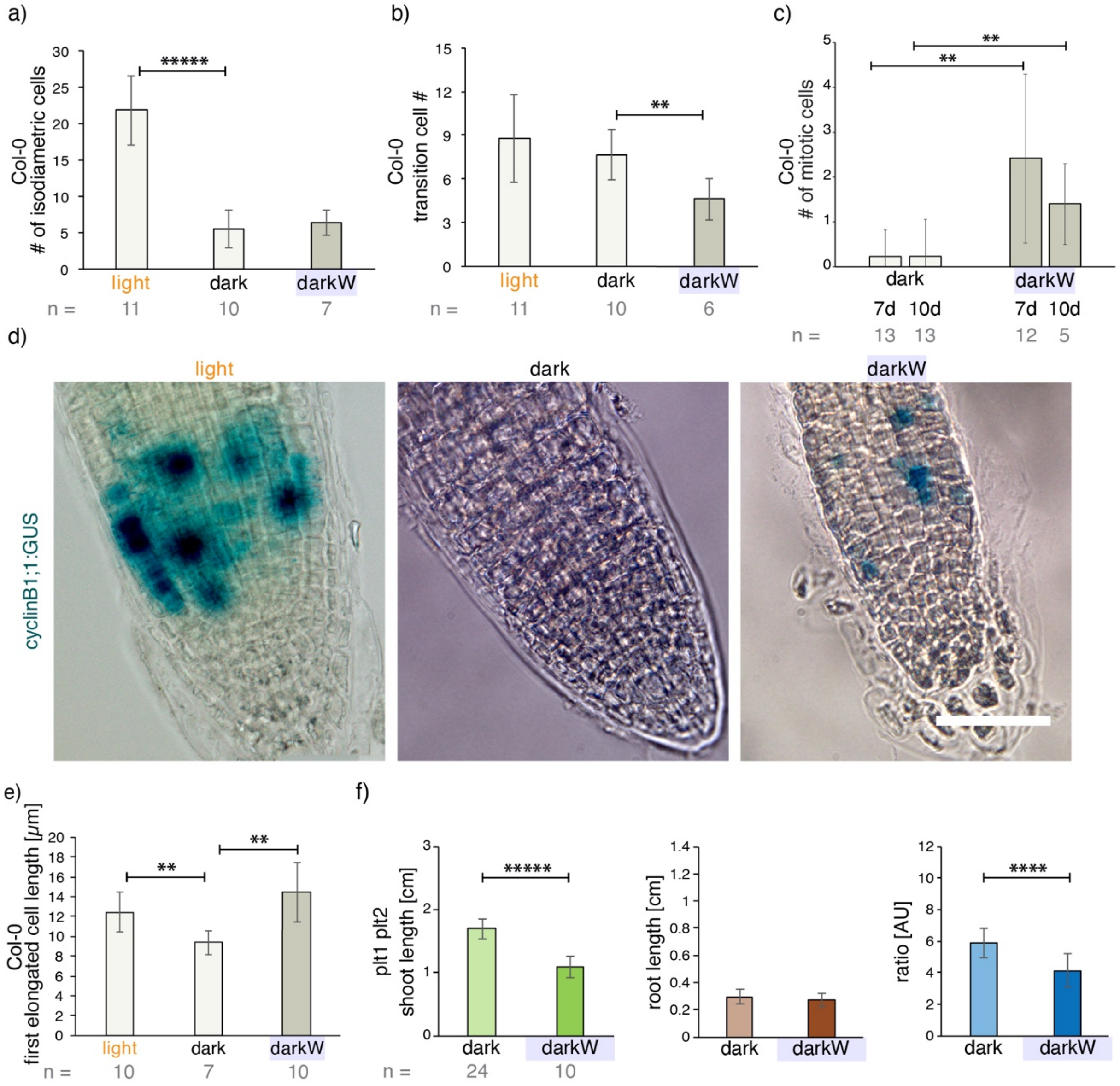


**Fig. S15 Root apical meristem properties under different environmental conditions.** a) A shift from light to darkness decreases the number of isodiametric cells (defined as being not longer than wide; $P = 1.2E^{-08}$) and a concomitant decrease in meristem size. (b) In the dark, water stress results in a decrease in the number of transition cells (includes the first cell being longer than wide up to the last cell whose length is <150% that of the previous cell; see Method S8). (c; d) Seedlings expressing the M-phase CycB:GUS marker and stained with X-Gal at day 7 (d) and at days 7 versus 10 (c). The number of cells expressing CycB:GUS cells per root tip decreased from light to dark (d) but increased from dark to darkW (c, d) at both days 7 and 10. (e) The The first elongated cell was longer under darkW (see orange asterisk in Fig. 7b) than under dark conditions. (f) *plt 1plt2* mutants have an unimpaired hypocotyl response but fail to reproducibly elongate their roots in response to water stress in the dark ($P_{root}$ = 0.27). The number (n) of seedlings measured per condition is in grey below the mean ±StDev bar graphs. *P*-values were computed with a non-parametric Mann-Whitney-U tests with a Benjamini–Hochberg correction in (c) and with a two-tailed student's *T*-test in panels a, b, e,f; they are represented as follows: *: 0.05 - 0.01; **: 0.01 - 0.001; ****: 0.0001 - 0.00001; *****: < 0.00001. Scale bars: 50 µm.

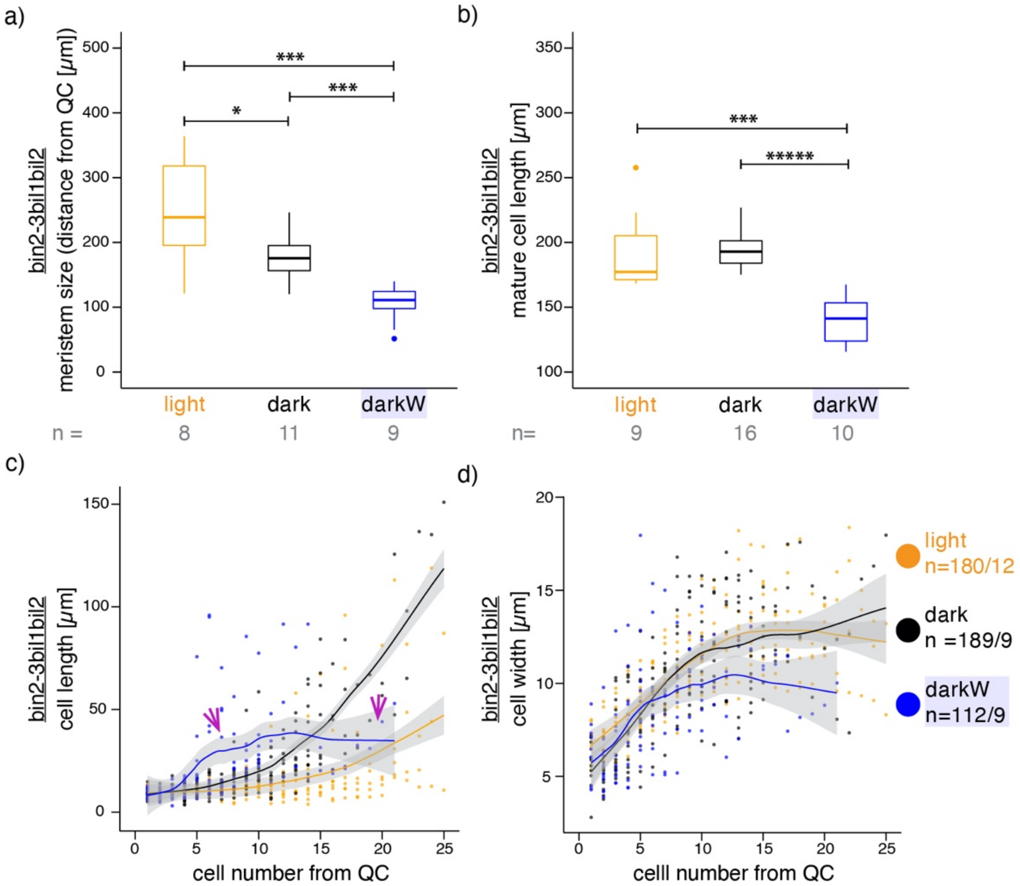


**Fig. S16. Root meristem properties in *bin2-3bil1bil2* under multiple stress conditions.** *bin2-3bil1bil2* seed were germinated in the light (orange), dark (black) or dark with -0.4MPa water stress (blue). 10 days after incubation, single epidermal cell files were measured, starting at the epidermal/ lateral root cap initials. (a) Meristem size determined via mixed Gaussian models, as described (Fridman et al. 2021; see Fig. S14). (b) Mature cell length, based on the ten most elongated cells for each condition. (c, d) Cell lengths (c) and width (d) of consecutive cells as a function of cell number from the quiescent center (QC); the fitted lines were generated with Local Polynomial Regression Fitting with the 'loess' method in R and grey shading designates the 95 percent confidence interval. The purple arrows point to the relatively flat slope for cell length (cf. green arrows pointing to the steep slope characteristic of the wild-type in Fig. 6d) in the darkW condition (c). The sample size (n) is given as the number of seedlings in panels a and b and as the number of cells/ number of seedlings that were analyzed in panels c, d. P-values were computed with a two-tailed student's T-test and are represented as follows: *: 0.05 - 0.01; ***: 0.001 - 0.0001; *****: < 0.00001. Related to Fig. 6.

**Table S1** Lines used in this study. The nature of the mutant alleles is depicted with null in regular font, semi-dominant or dominant in bold and higher order mutants underlined.

| Allele | NASC accession | AGI | ecotype | dominance | Reference |
|---|---|---|---|---|---|
| *cry1-1 cry2 phyA-201 phyB-5* | | At4g08920 At1g04400 At1g09570 At2g18790 | Ler | quadruple null | Mazella and Casal, 2001 |
| *det2-1* | N6159 | At2g38050 | Col-0 | null | Chory et al., 1991 |
| *Cpd* | N511386 | At5g05690 | Col-0 | null | Szekeres et al., 1996 |
| *bri1-6* | N399 | At4g39400 | En-2 | hypomorphic | Noguchi et al., 1999 |
| *bri1-116brl1brl3* | | At4g39400, At1g55610, At3g13380 | Col-0 | triple null | Kang et al., 2017 |
| *bak1-1* | N6125 | At4g33430 | Ws-2 | null | Li et al., 2002 |
| ***bin2-1*** | | At4g18710 | Col-0 | **semi-dominant** | Li et al., 2001 |
| ***ucu1*** | | At4g18710 | Ler | **semi-dominant** | Pérez-Pérez et al., 2002 |
| ***dwarf12*** | | At4g18710 | Ws-2 | **semi-dominant** | Choe et al., 2002 |
| *bin2-3bil1bil2* | | At4g18710, At2g30980 At1g06390 | Ws-2 | triple null | Yan et al., 2009 |
| ***bzr1-1D*** | N65987 | At1g75080 | Col-0 | **dominant** | Wang et al., 2002 |
| ***bes1-D*** | N65988 | At1g19350 | En-2 | **semi-dominant** | Yin et al., 2002 |
| *bri1-116* ***bzr1-1D*** | | At4g39400, At1g75080 | Col-0 | null, dominant | Wang et al., 2002 |
| *plt1plt2* | | At3g20840 At1g51190 | WS | double null | Aida et al., 2004 |

**Table S2** Segregation analysis of the B1 mapping population. Of 19 F2 individuals sequenced at the BIN2 TREE domain, there was an absolute segregation between the sequence and the phenotype. Thus, plants homozygous for the wild-type TREE domain had a wild-type phenotype, TREE/TREK plants heterozygous for the TREE domain had an intermediate phenotype (clear BR phenotype with rolled in leaves but medium stature and fertile) and plants homozygous for the B1 TREK mutation were semi sterile dwarfs with severe BR phenotypes. Related to Fig. 2.

| F2 line # | BIN2 TREE domain | Phenotype |
|---|---|---|
| 21 | heterozygous TREE/TRKE | Intermediate |
| 22 | homozygous TREE | wild-type |
| 23 | homozygous TREE | wild-type |
| 24 | homozygous TREE | wild-type |
| 25 | heterozygous TREE/TRKE | Intermediate |
| 26 | heterozygous TREE/TRKE | Intermediate |
| 27 | homozygous TREE | wild-type |
| 28 | heterozygous TREE/TRKE | Intermediate |
| 29 | heterozygous TREE/TRKE | Intermediate |
| 30 | heterozygous TREE/TRKE | Intermediate |
| 31 | homozygous TREE | wild-type |
| 32 | heterozygous TREE/TRKE | Intermediate |
| 33 | homozygous TREE | wild-type |
| 34 | homozygous TREK | Mutant |
| 35 | homozygous TREK | Mutant |
| 40 | homozygous TREK | Mutant |
| 41 | homozygous TREK | Mutant |
| 42 | homozygous TREK | Mutant |
| 45 | heterozygous TREE/TRKE | Intermediate |

**Method S1 Composition of nutrient stress plates**

The following macronutrients were used for nutrient stress conditions:

| | NPK | -P | -N | -K | MS |
|---|---|---|---|---|---|
| KNO3 | 5 mM | 5 mM | 0 mM | 0 mM | 19 mM |
| Ca(NO3)2 | 2 mM | 2 mM | | 2 mM | |
| MgSO4.7H2O | 2 mM | 2 mM | 2 mM | 2 mM | 1,5 mM |
| KH2PO4 | 2,5 mM | | 2,5 mM | | 1,25 mM |
| KCl | | 2,5 mM | 5 mM | | |
| (NH4) 2HPO4 | | | | 2,5 mM | |
| CaSO4 | | | 2 mM | | |
| (NH4) NO3 | | | | | 20,6 mM |
| CaCl2 | | | | | 2,23 mM |

**Macronutrients used for nutrient stress or MS.**

Media were solidified with 1.2 % (w/v) agar.

**Method S2 Supplemental information on light experiments**

Light intensities were determined with spectroradiometers (white, blue, and red light: model Li-1800 [LiCor]; far-red light: model SKP200 with a sensor for 730 nm [Skye Instruments]). The blue, red, and far-red light sources were generated by light-emitting diodes using diodes with emission maxima at 469, 660, and 740 nm (Quantum Devices; PVP).

**Method S3 Preparation of PEG plates**

Square plates (120 mm x 120 mm) were poured with exactly 45 mL (½x or 1x) MS medium. The 2x PEG-solution was prepared according to the following table:

| water potential (Mpa) | MS | 2x PEG-6000 (g/L) |
|---|---|---|
| 0 | 1/2 x | 0 |
| -0.2 | 1x | 0 |
| -0.3 | 1/2 x | 220 |
| -0.4 | 1/2 x | 280 |
| -0.5 | 1/2 x | 320 |
| -0.6 | 1x | 280 |
| -0.7 | 1x | 320 |

**MS strength (1/2x, 1x) and PEG concentrations used for a gradient of water stress.**

It is to be noted that there was some variability between different PEG-6000 lots from the same provider, such that each lot needed calibration. The 2x PEG solution was sterile filtered and exactly 45 mL were added onto polymerised plates. After an incubation of exactly 24 hours at RT, the PEG-solution was completely decanted.

Plastic strips 2.5 cm x 10 cm were cut from transparencies, making sure the cuts were straight. The strips were twice sterilised in 80 % EtOH for 5 min and air dried. Two strips were placed on each plate as follows: the strip was carefully pushed 1 mm deep into the agar at max. 30°. Then, the strip was leaned onto the agar, making sure the agar was not damaged in the process.

**Method S4 Seed handling for screen**

Fresh (max. two-year-old, stored at RT) or frozen (-20 °C) seed stocks were used from plants grown under optimal chamber conditions. Seed were surface sterilised using a brief 80% ethanol rinse followed by 15 min incubation in sterilisation buffer (0.01 % SDS, 3% NaOCl). After 5 washes in mQ water, seed were resuspended in 0.15 % agar and imbibed in the dark at 4 °C for 7 days to break dormancy. Seed were pipetted at the interface between the foil and agar such that, upon germination, only the root touched the agar. Mutants were sown on the same plates as the corresponding wild type ecotype (Table S1). For water stress, seeds in each biological replicate were sown on 2 plates and these then pooled for the analysis. Plates were sealed with breathable (or porous) tape (https://www.soehngen.com). Plates for dark conditions were wrapped with two layers of thick aluminium foil. All plates were negatively inclined by 4° to promote root growth on the surface rather than in the agar. Incubation was for 10 days at 22 °C in a growth chamber with a permanent light (180 µmol $m^{-2}s^{-1}$).

**Method S5 Picking, scanning, phenotyping and genotyping.**

10-day old seedlings were transferred onto cold 1.2 % Agar plates and scanned at 1200 ppi. The images were saved as tiff files. Pixel measurements on hypocotyl and root were performed in Fiji-ImageJ using the free-hand-tool. The ratio was calculated as hypocotyl/root. Light versus dark comparisons were computed as light/dark and dark versus darkW comparisons as dark/darkW. Threshholds are described in Fig. S7 for dark versus darkW and in Fig. S9 light versus dark. Where possible, we avoided segregating lines. For *bin2-1*, we were able to obtain homozygous *bin2-1* lines by propagating plants under optimal growth conditions over > 4 months at the TUMmesa ecotron. We distinguish homo- and heterozygous seed by (i) adult phenotype (ii) segregation in the next generation (iii) root:hypocotyl ratios from dark-grown seedlings on plate and (iv) sequencing of the TREE domain (as shown in Fig. 2e). For the non-viable segregating lines *cpd* and *bri1brl1brl3* lines, which have a null *bri1-116* allele and segregate (*bri1-/+ brl1-/- brl3 -/-*), phenotypes were verified by propagating seedlings on plate after the scan on day 10 and scored when the button like rosette and dwarf phenotypes were clearly apparent.

**Method S6 Positional Cloning**.

Segregation analysis was carried out on 345 F2 individuals with mapping primers closely linked to the B1 mutation (see Fig. 2). Segregation analysis established that (i) the mutation was semidominant and (ii) the phenotype segregated with the closely linked markers on chromosome 4. 19 F2 plants were subsequently sequenced at the BIN2 TREE domain, as shown in Table S2. The genomic BIN2 TREE domain was amplified using the Phire Plant Direct PCR Kit (Thermo Fischer Scientific) according to the manufacturer's instructions. The primers used for the amplification PCR were

bin2_TREE forward CAC CCG AGC TCA TAT TTG GT
bin2_TREE reverse CTT CTG GGG GCA TCC TTT TG.

Annealing was at 60 °C for 5 s. The BIN2 PCR products were analyzed by Sanger sequencing with the forward primer. Wild-type plants as well as homozygous and heterozygous *bin2-1* mutants were distinguished based on the sequencing chromatograms. The wild type and homozygous *bin2-1* had a single peak in the sequencing trace at position 989, coding for G or A, respectively. Instead, heterozygous *bin2-1* mutants showed a twin pair of G and A peaks for the TREE domain at this base position.

| marker name | Mb | indel/ enzyme | forward primer (5'->3') | reverse primer (3'->5' | Col (bp) | Ler (bp) | annealing |
|---|---|---|---|---|---|---|---|
| CER453988 | 9,85 | 45/-45 | Tcccctaagcccatacatcg | Ttatgcatcgtcggttggtag | 280 | 235 | 55 °C 30 s |
| CER466027 | 10,01 | DdeI | Catcaatattcggcactcca | gttgtgacatcggctccttt | 287+114 | 400 | 52 °C 30 s |
| CER451720 | 10,15 | 19/-19 | gattggttggttttcccattt | Caaacgcagcaacatcagtt | 245 | 225 | 52 °C 30 s |
| CER430180 | 10,22 | TaqI | cgattcatgtttccgtgatg | Aaacaggtgccatcatttcc | 387+202 | 587 | 52 °C 30 s |
| CER451656 | 10,28 | 29/-29 | Aagcacattcaaacaaaatctcc | Agcggaaaattctgatggtg | 314 | 285 | 52 °C 30 s |
| CER465938 | 10,39 | Hpy188I | tccatcctcctcctccttct | Cctggatggaaatggatgtt | 226+108 | 33 | 53 °C 30 s |
| CER446965 | 10,59 | Bsu15I | Agattcgccaggttatccaa | ttcggatcgatcatttttcc | 96+496+11 | 599 | 50 °C 30 s |
| CER466234 | 10,85 | 58/-58 | tctctatttccggcgactgt | Ccgtcacaatcctgactcaa | 300 | 242 | 55 °C 30 s |

**Markers and primers used for positional cloning on chromosome 4.**

**Method S7 Real-time PCR analysis**

Total RNA was extracted with the RNeasy Plant Mini Kit (Qiagen) according to the manufacturer's protocol. A 0.5 µg aliquot of RNA was subjected to first-strand cDNA synthesis using iScript™ cDNA Synthesis Kit (Bio-Rad) according to the manufacturer's protocol. The cDNA was diluted 50-fold. The gene-specific primers used for real-time PCR were:

| | | forward | Reverse |
|---|---|---|---|
| *LHCB1.2* | At1g29910 | CCG TGA GCT AGA AGT TAT CC | GTT TCC CAA GTA ATC GAG TC |
| Ubiquitin-protein ligase-like protein | AT4G36800 | CTG TTC ACG GAA CCC AAT TC | GGA AAA AGG TCT GAC CGA CA |

Real-time PCR analysis was performed using a BioRad Real-Time System CFX96TM C1000 Thermal Cycler using a SsoAdvanced Universal SYBR Green Supermix (BioRad). Relative transcript abundance of *LHCB1.2* (*LIGHT HARVESTING CHLOROPHYLL A/B BINDING PROTEIN 1.2*) was normalized with respect to the level of the constitutively expressed mRNA for Ubiquitin-protein ligase-like protein using the Bio-Rad CFX Maestro Software. Melting curve analysis was performed to check for non-specific PCR products and primer dimers. For each sample each experiment was repeated at least three times with three technical replicates.

**Method S8 confocal microscopy and root apical meristem properties**

Confocal microscopes used for imaging were an Olympus (www.olympus-ims.com) Fluoview 1000 confocal laser scanning microscope (CSLM) and a Leica (www.leica-microsystems.com) SP8 Hyvolution CSLM. 40x and 60x water immersion 0.9 numerical aperture objectives (Olympus) were used. For FM4-64 staining emission was at 640 nm. Six-day-old Col-0 seedlings were used for counting isodiametric and transition cells and 10-day old seedlings were used for mixed Gaussian model analysis (Fig. S14; Fridman et al., 2021). For cell counting, epidermal cells in the meristematic zone were divided into isodiametric versus transitioning cells; isodiametric cells were counted including the first cell diagonal from the QC (epidermis initial) and the last cell not being longer than wide. Transitioning cells include all cells from the first cell being longer than wide to the last cell having less than 150% the length of the previous one; cells in the elongation zone had twice the length of the immediately preceding cell (González-García et al., 2011).

**Methods S9 GUS staining**

Seed were plated under initial screen conditions (MS salts) and 7- or 10-day-old seedlings were used for GUS staining. Seedlings were fixed for 20 min in 90 % acetone, washed with staining buffer (10 mM EDTA, 0.1 % Triton-X 100, 2 mM potassium ferrocyanide, 2mM potassium ferricyanide, 50 mM sodium-phosphate buffer (pH 7)) on ice and transferred into staining buffer containing additionally X-Gluc (2.31 mM) and chloramphenicol (100 mg/ml). Seedlings were incubated at 400 mbar for 15 min on ice before being incubated for 3-5 h at 37 °C. Seedlings were dehydrated in an ethanol series (20 %, 35 %, 50 %, 30 min per concentration) and finally fixed for 30 min at room temperature in FAA (50 % ethanol, 3.7 % formaldehyde, 5 % acetic acid). Micrographs were recorded with an Olympus BX61 microscope. A 40x objective (water) was used.

*Supplementary references*

Chory, J.; Nagpal, P.; Peto, C. A. (1991): Phenotypic and Genetic Analysis of det2, a New Mutant That Affects Light-Regulated Seedling Development in Arabidopsis. In: *The Plant Cell* 3 (5), S. 445–459. DOI: 10.1105/tpc.3.5.445.

Noguchi, T.; Fujioka, S.; Choe, S.; Takatsuto, S.; Yoshida, S.; Yuan, H. et al. (1999): Brassinosteroid-insensitive dwarf mutants of Arabidopsis accumulate brassinosteroids. In: *Plant physiology* 121 (3), S. 743–752. DOI: 10.1104/pp.121.3.743.
